# Residue retention promotes soil carbon accumulation in minimum tillage systems: Implications for conservation tillage

**DOI:** 10.1101/746354

**Authors:** Yuan Li, Zhou Li, Scott X. Chang, Song Cui, Sindhu Jagadamma, Qingping Zhang, Yanjiang Cai

## Abstract

Crop residue retention and minimum tillage (including no-tillage, NT, and reduced tillage, RT) are common conservation tillage practices that have been extensively practised for improving soil health and reducing the negative environmental impact caused by intensive farming. However, the complex effect of conservation tillage practices on soil organic carbon (SOC) storage has not been systematically analyzed, and particularly, the synergistic effect of crop residue retention and minimum tillage on SOC storage remains nonexistent. We conducted a global meta-analysis using a dataset consisting of 823 pairs of data points from 164 studies. We analyzed the effect of crop residue retention and minimum tillage on SOC storage and how the above effects were influenced by various soil/environmental (soil sampling depth, soil texture, and climate) and management conditions (cropping intensity and treatment duration). We found that either residue retention or minimum tillage alone increased SOC stock, while the former increased SOC more. The NT and RT increased SOC stock by 10 and 6%, respectively, in comparison to conventional tillage (CT). The NT plus residue retention (NTS) and RT plus residue retention (RTS) resulted in 20 and 26% more increase in SOC than NT and RT, respectively. Compared with CT, NTS and RTS further increased SOC stock by 29 and 27%, respectively. The above effects were greater in the topsoil than in the subsoil. Availability of initial soil nutrient played a greater role in affecting SOC stock than climatic conditions and management practices. Both residue retention and NT increased SOC rapidly in the first 6 years regardless of soil texture or climate condition, followed by a period of slower sequestration phase before reaching a slow steady rate. Double cropping generally increased SOC stock across all conservation tillage practices as compared to single or multiple cropping. Therefore, we conclude that minimum tillage coupled with residue retention in a double cropping system is the most beneficial management system for increasing cropland SOC storage, which can inform sustainable soil management practices aimed at increasing global C sequestration.

## 1. Introduction

Nearly 1.3% of the global land area is dedicated for crop production (FAO, 2015), while intensification of crop production has contributed a large amount of greenhouse gases (such as carbon dioxide-CO_2_, nitrous oxide-N_2_O, and methane-CH_4_) to the atmosphere (Strassmann et al., 2008). It is well acknowledged that soil organic carbon (SOC) in croplands is the core of soil fertility that ensures crop production and food security, but the loss of C from cropland through land cultivation has fundamentally altered the global C cycle (Lal, 2004b; Smith et al., 2005). The C loss is mainly caused by the post-harvest removal/loss of crop residues and C distribution/decomposition processes from agricultural fields, such as hay harvest or tillage (Lal, 2004a). Since SOC sequestration is crucial for soil fertility and climate change mitigation, alternative production practices have been adopted globally to enhance soil C storage (Chenu et al., 2019; Lal, 2004b; Sauvadet et al., 2018).

Conservation tillage practices, a term originally introduced by Wall (2006), encompass three managerial components, i.e., minimum tillage (including no-tillage, NT, and reduced tillage, RT) with or without crop residue retention. It has been estimated that a SOC sequestration rate of 0.4-0.8 Pg C yr^-1^ can be achieved through the adoption of conservation tillage practices across all cropping systems globally, which may account for 33-100% of total SOC sequestration potential in the world (Lal, 2004a). Furthermore, converting from conventional tillage (plough tillage, CT) to NT alone can potentially increase SOC at a rate of 0.1-1 Mg C ha^−1^ yr^−1^ (Lal, 2004a). In addition, NT can improve soil structural stability, soil nutrient availability, and water holding capacity (Blanco-Canqui and Ruis, 2018; Schmidt et al., 2018), which eventually lead to greater crop productivity and SOC sequestration. However, a global analysis in 2010 indicated that NT benefits SOC only in the top 10 cm (Luo et al., 2010). Furthermore, Follett et al. (2013) found no difference in SOC in the 0-120 cm soil profile between NT and CT after 8 years in an irrigated corn (*Zea mays*) field. Based on data from 62 studies, similar results were also observed in moist soils of eastern Canada (VandenBygaart et al., 2003). In addition, residue retention can directly increase C input into the soil, as well as improve soil structure and nutrient availabilities, and increase soil microbial population size (Johnson and Hoyt, 1999; Li et al., 2018). The increased cropping intensity can increase the production of both above- and belowground biomass that can be later incorporated into the overall SOC pool (Luo et al., 2010). However, the increment of SOC stock in response to a combination of conservation tillage practices has not been quantitatively evaluated. Quantitative analyses have only been conducted on individual conservation tillage practices measures to assess the effect of NT (Luo et al., 2010), straw retention (Han et al., 2018; Lu et al., 2009), or cover crops (Poeplau and Don, 2015) on SOC storage. Those quantitative analyses have found great variations in responses under individual conservation tillage practices. For instance, straw retention alone sequestered more SOC than NT alone in China (Lu et al., 2009), and the effect of NT on sequestrating C was strongly regulated by cropping systems (Luo et al., 2010). Other factors, such as study duration, climate condition, and soil texture, were also critical in affecting the direction and magnitude of changes in SOC storage in response to conservation tillage practices (Chenu et al., 2019; González-Sánchez et al., 2012; Lal, 2004a; Luo et al., 2010).

As mentioned earlier, it can be concluded that even though the agronomic and ecological benefits associated with different conservation tillage practices have been widely recognized, the combined effects of multiple conservation tillage practices have not been quantitatively evaluated. This knowledge gap impedes our ability to implement site-specific conservation tillage practices that can enhance SOC reserve and prevents researchers from gaining a deeper understanding on the synergistic effects of different conservation tillage practices and their combined effects as a whole in different agroecosystems. In this meta-analysis, we hypothesized that residue retention coupled with minimum tillage sequester more SOC than residue retention or minimum tillage alone. Limited by data availability/quality and the scope and complexity of statistical inferences, we primarily focused on two aspects of the conservation tillage practices, residue retention and tillage intensity, while cropping intensity was treated as an additional management factor. Our objectives were to 1) determine the direction and magnitude of change in SOC stock in response to different combinations of residue retention and tillage practices; and 2) evaluate the effect of external abiotic (e.g., soil texture, climate factors) and management practices (cropping intensity, duration of conservation tillage practices) on SOC stock.

## 2. Materials and methods

### 2.1 Data collection

Published journal articles from January 1987 to April 2018 were searched using the ISI Web of Science (http://apps.webofknowledge.com/) and China Knowledge Resource Integrated Database (http://www.cnki.net/) using the keywords: C stock, C sequestration, soil quality, chemical properties, conservation practice, tillage, till. To minimize bias, the following criteria were followed: (1) paired observations involving a CT - CT with (CTS) or without residue retention (CT), and only field-based experiments were included. Specific conservation tillage practices chosen for comparison were RT or RT plus residue retention (RTS), and NT or NT plus residue retention (NTS). If a study used more than one conservation tillage practice at the same site, different conservation tillage practices were separately paired with the CT control; (2) the mean and standard deviation (SD) or standard error (SE) of the results obtained from replicated field plots (> two replications) were used in the meta-analysis; (3) the residue retention and tillage practices were treated as main treatments, while keeping other agronomic practices and factors such as cropping intensity, study duration, fertilizer management, and irrigation the same among the main treatments.

In total, 164 peer-reviewed publications covering all continents were included (Fig. 1). Soil bulk density (BD), and soil C and nitrogen (N) stock data were directly obtained from tables and texts of articles, or extracted from figures using the Get-Data Graph Digitizer software (ver. 2.24, Russian Federation). The final dataset included 1944 data points for soil C concentration, 996 data points for soil N concentration, and 1607 data points for soil BD under different conservation tillage practices. Details of the selected studies and references were presented in the supplementary material. Particularly, in certain countries such as the US, crop residues are typically retained in the field after harvest regardless of tillage practices, which were only considered as NTS, RTS, or CTS in this study.

**Figure 1.**
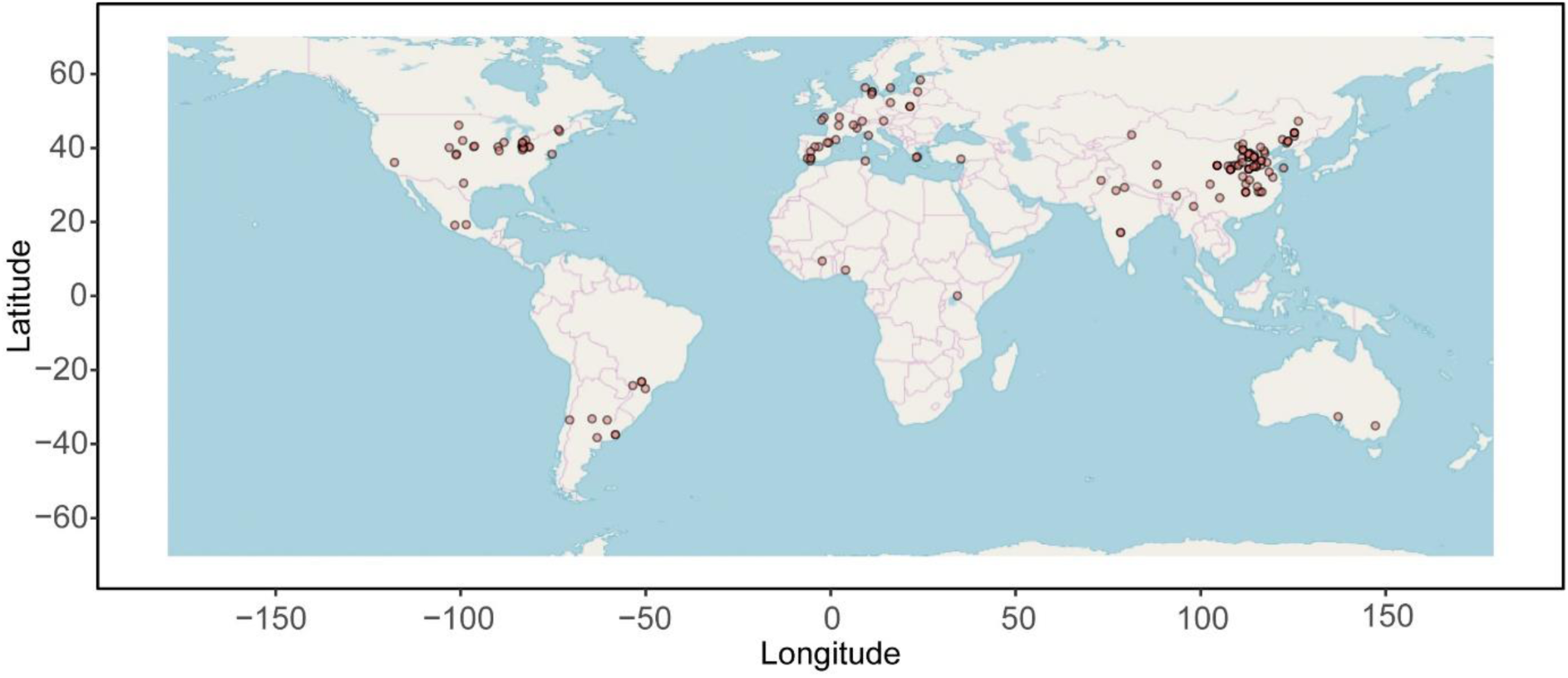
The geographical coverage of the studies used in the meta-analysis (the colour version can be found in the web version of this article).

For each site, we also extracted information about experimental and management factors (e.g., duration of the experiment, climatic parameters, cropping intensity), and soil properties (e.g., soil texture, soil pH, and initial C and N concentrations). The data were then grouped by (i) soil texture, i.e., sandy (sand, loamy sand, and sandy loam), silty (silty loam and silt), loamy (sandy clay loam, loam, clay loam, and silty clay loam), or clayey (clay, sandy clay, and silty clay); (ii) the duration of the study (<6, 6–12, >12 years); (iii) cropping intensity (single-, double- and multi-cropping in a given year); (iv) climatic parameters (mean annual temperature, MAT, and mean annual precipitation, MAP); and (v) soil sampling depth (topsoil, 0-15 cm, and subsoil, >15 cm). In particular, MAT values were used for generating the aridity index (AI), which was calculated from the equation AI = MAP/(MAT + 10). The AI values of 0-10, 10-20, 20-30, and >30 correspond to arid or semi-arid, semi-humid, humid, and perhumid environments, respectively (De Martonne, 1926). However, due to restrictions caused by limited data (Table S1), N stock was not selected for subgroup meta-analysis according to experimental and management factors. Additionally, the comparison of RTS *vs.* RT, was excluded in subgroup meta-analysis due to insufficient data; only semi-humid, humid, and perhumid climates were studied due to insufficient data from arid or semi-arid regions.

### 2.2 Data analysis

For studies that did not report soil C or N stock values, we used both soil BD and C or total N (STN) concentration to calculate the C or N stock: SOC (or STN) stock = C_SOC_ (or C_STN_) × BD × D, where C_SOC_ and C_STN_ is the SOC and STN concentration (g kg^-1^), respectively; BD is bulk density (g cm^-3^) of the corresponding soil layer and D is the thickness of the soil layer (cm) (Lal, 2004a). The natural logarithm of the response ratio (ln*RR*) of soil C or N stock (Mg ha^-1^) between treatment and control was computed according to Osenberg et al. (1999) and following methods used in Li et al. (2018). The variance (*v*) of the ln*RR* was calculated from the equation, *v* = S_t_ /n_t_ × X_t_ + S_c_ /n_c_ × X_c_, where S_t_ and S_c_ represent the standard deviations of the treatment and control groups, and n_t_ and n_c_ represent the number of replicates of the treatment and the control groups, respectively. For studies that did not report either SD or SE, SD was estimated as 0.1 times the mean (Luo et al., 2006). To derive the overall response effect of the treatment group relative to the control group, the weighted response ratio (*RR*_++_) between treatment and control groups was calculated according to Hedges et al. (1999); Luo et al. (2006), as described in Equation (1):

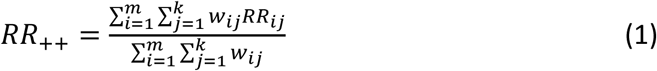

Where *m* represents the number of groups compared, *k* represents the number of comparisons in the corresponding group, and *w_ij_* represents the weighting factor (*w_iJ_* = 1/ *v_ij_*). The standard error of *RR*_++_ was calculated using Equation (2):

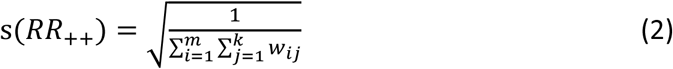

The 95% confidence interval (95% CI) was calculated by *RR*_++_ ± 1.96 × s(*RR*_++_). When the 95% CI value of the response variable did not overlap with 0, the conservation tillage practice effect on the variable was considered to be significantly different between the control and the treatment groups (Gurevitch and Hedges, 2001). Additionally, the percentage change was calculated as RR_++_: (exp(*RR*_++_) - 1) × 100%.

The meta-analysis was performed using the ‘metafor’ package (Viechtbauer, 2010) in the R statistical software (version 3.4.2). For each group (soil texture, study duration, cropping intensity, AI, soil sampling depth), the mean effect size (ln*RR*) and its 95% CI were calculated with bias-correction generated via bootstrapping. A Chi-square test was used to determine whether the heterogeneity among the ln*RR* of changes in SOC and N stocks under conservation tillage treatments (Q_total_) significantly exceeded the expected sampling error. A randomization test was used to determine the significance of the between-group heterogeneity (Q_b_) (Adams et al., 1997). Chi-square tests were also used to determine whether the remaining within-group heterogeneity (Q_w_) was significant (see Table S2-S7).

The effect of environmental and management factors (independent variables) on the SOC stock was determined using linear regression analyses. Correlations between independent variables were detected using the Kendall correlation analysis. For categorical variable, soil sampling depth (topsoil or subsoil) was converted to binary data, and cropping intensity was converted to numeric data by counting the number of crops (number of crops from the same field during one agricultural year) prior to conducting the analysis. The network plot of the correlation of C stock to soil properties, environmental and management conditions, and the correlation data frame were conducted and plotted through the “corrr” package (Jackson, 2016) in R (version 3.4.2). The identified variables having a stronger correlation coefficient with C stock were further used to make regression analyses to quantify the effects of those variables on C stock. The associated regression analyses and plots were made using the “ggplot2” package (Wickham, 2016).

## 3. Results

### 3.1 Effect of conservation tillage practices on C and N stocks

Relative to CT, minimum tillage (including NT and RT) increased SOC stock (Fig. 2a, *p* < 0.01), and NT increased SOC stock 10% more than RT (*p* < 0.001, percentage change was calculated based on the RR_++_). Residue retention significantly increased SOC stock regardless of tillage intensity, RTS, NTS and CTS resulted in 26, 20, and 22% greater SOC stock than RT, NT, and CT, respectively. While no difference was observed between NTS and RTS (*p* = 0.08). The NTS and RTS significantly increased SOC stock by 29 and 17%, respectively, in comparison with CT. Minimum tillage also significantly increased the soil N stock (Fig. 2b). Residue retention did not lead to a significant increase in soil N stock, with the exception of NTS as compared with NT (*p* = 0.03). Additionally, the interaction between minimum tillage and residue retention also did not affect soil N stock.

**Figure 2.**
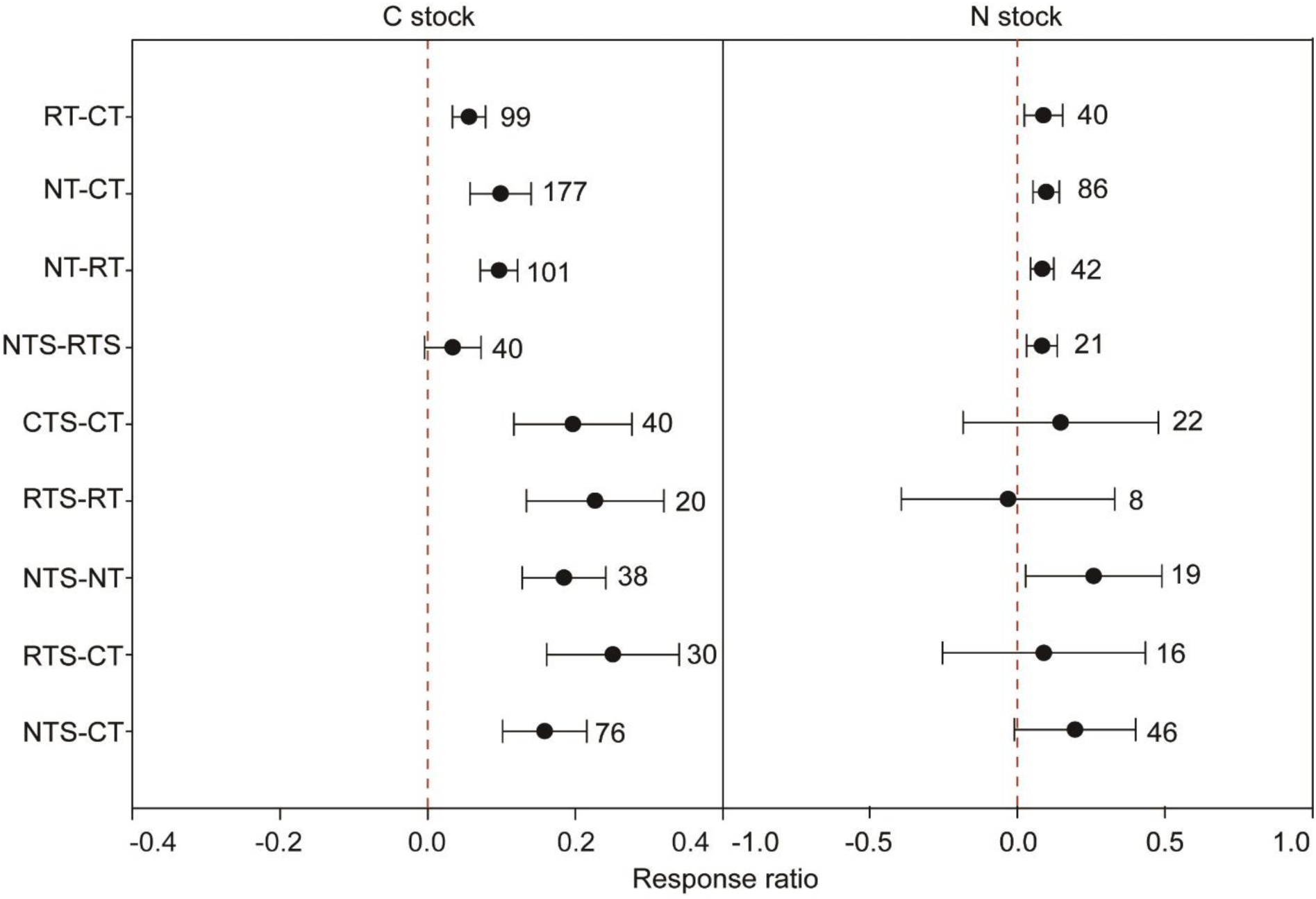
The effect size for a) soil organic carbon (C) stock, and b) nitrogen (N) stock under different conservation agricultural practices. Effect size stands for the weighted response ratio between treatment and control. Error bars represent the 95% confidence intervals. The sample size of each variable is noted adjacent to each bar. The abbreviations are no-tillage (NT), reduced tillage (RT), and conventional tillage (CT) without residue retention and no-tillage (NTS), reduced tillage (RTS), and conventional with residue retention (CTS).

### 3.2 How do management conditions change the effect of conservation tillage practices on C stocks

The SOC stock increased under NT compared with CT and under RTS compared with CT under single cropping intensity condition (Fig. 3a, *p* < 0.001). However, NTS significantly decreased SOC stock as compared with NT with an effect size of −0.18. Under the double cropping system, changes in SOC stock were significant for all treatments (Fig. 3b). Under multiple cropping intensity conditions, only NTS and NT increased SOC stock as compared with NT and CT, respectively (Fig. 3c, *p* < 0.01). While the confidence interval for NTS *vs.* CT or CTS *vs.* CT was very wide and included zero, indicating that there was a high amount of variability remaining in the data.

**Figure 3.**
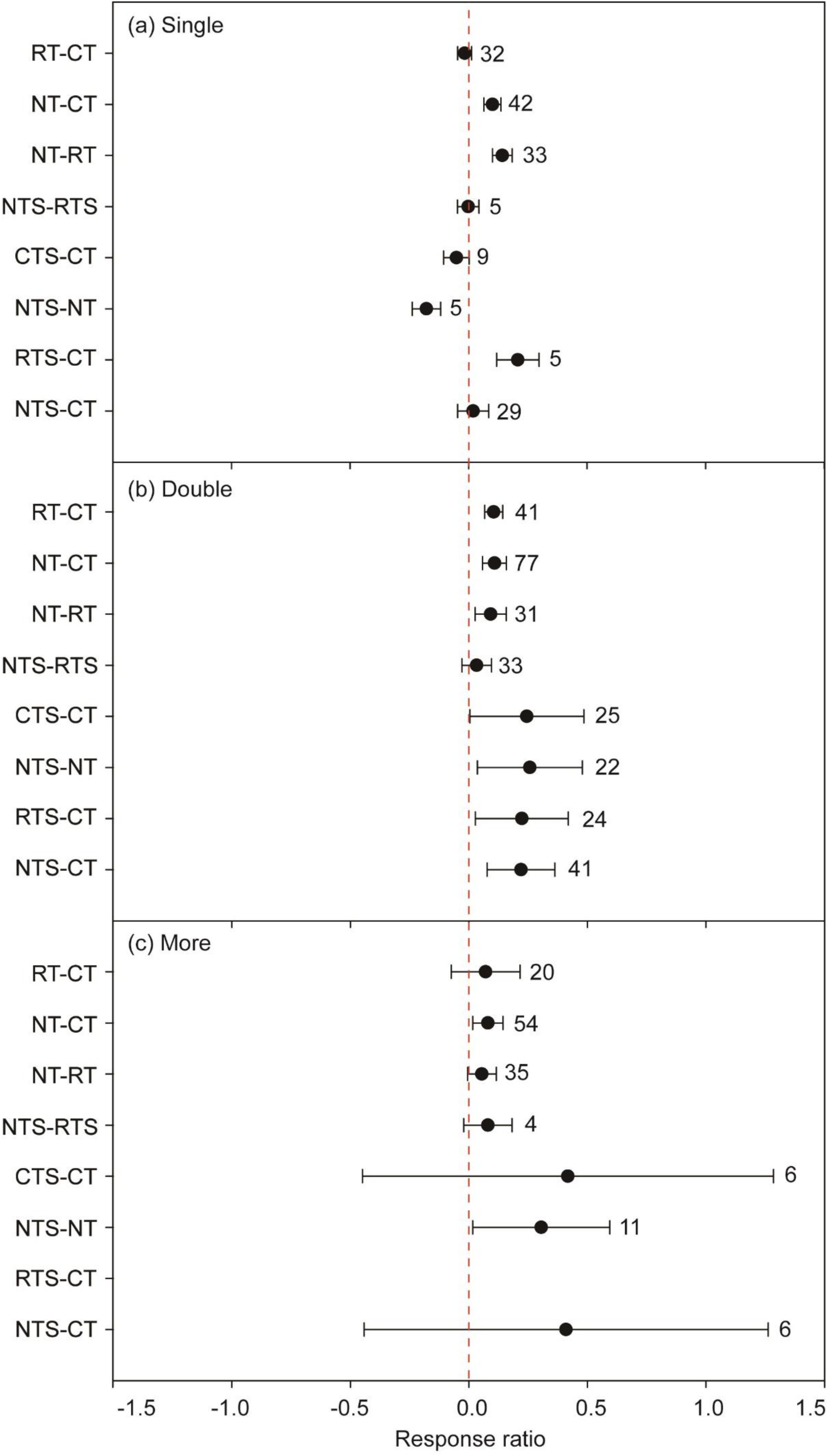
The effect size for soil organic carbon stock under different conservation agricultural practices among cropping intensity, a) single, b) double, and c) multiple. Effect size stands for the weighted response ratio between treatment and control. Error bars represent the 95% confidence intervals. The sample size of each variable is noted adjacent to each bar. For a description of the abbreviations, refer to the Fig. 2 title.

When the study duration was <6 years, NT increased SOC stock relative to CT (Fig. 4a, *p* < 0.001), and RTS but not RT significantly increased SOC stock in comparison to CT. Compared to CT, NT or CTS, NTS increased SOC stock (*p* < 0.001). The NTS resulted in 171% and 273% higher SOC stock than NT and CT by, respectively (*p* < 0.05). All conservation tillage practices significantly increased SOC stock as compared to CT when the study duration was between 6 and 12 years, with the exception of NT (Fig. 4b). In long-term (>12 years) studies, NT and RT significantly increased SOC stock as compared to CT (Fig. 4c), and the stock was significantly higher in NT than RT. However, CTS decreased SOC stock by 70% as compared to CT (*p* = 0.04), and NTS significantly decreased SOC stock by 48% as compared to NT in long-term studies.

**Figure 4.**
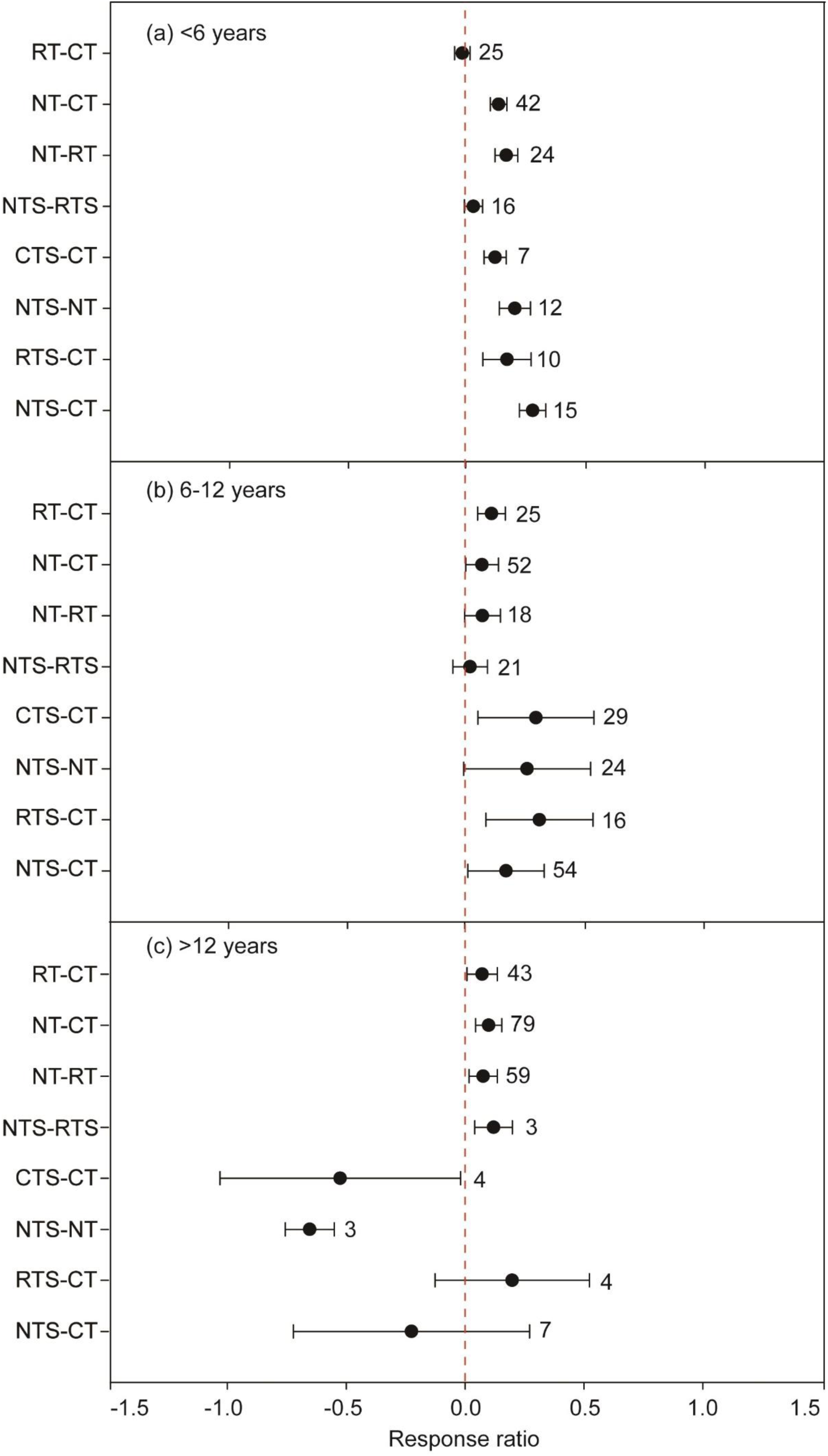
The effect size for soil organic carbon stock under different conservation agricultural practices over study durations, a) <6 years, b) <6-12 years, and c) >12 years. Effect size stands for the weighted response ratio between treatment and control. Error bars represent the 95% confidence intervals. The sample size of each variable is noted adjacent to each bar. For a description of the abbreviations, refer to the Fig. 2 title.

### 3.3 How do environmental conditions change the effect of conservation tillage practices on C stocks

Changes in SOC stock were significant for all comparisons in the topsoil (Fig. 5a). However, in the subsoil, NTS led to a significant decrease in SOC stock relative to CT (Fig. 5b).

**Figure 5.**
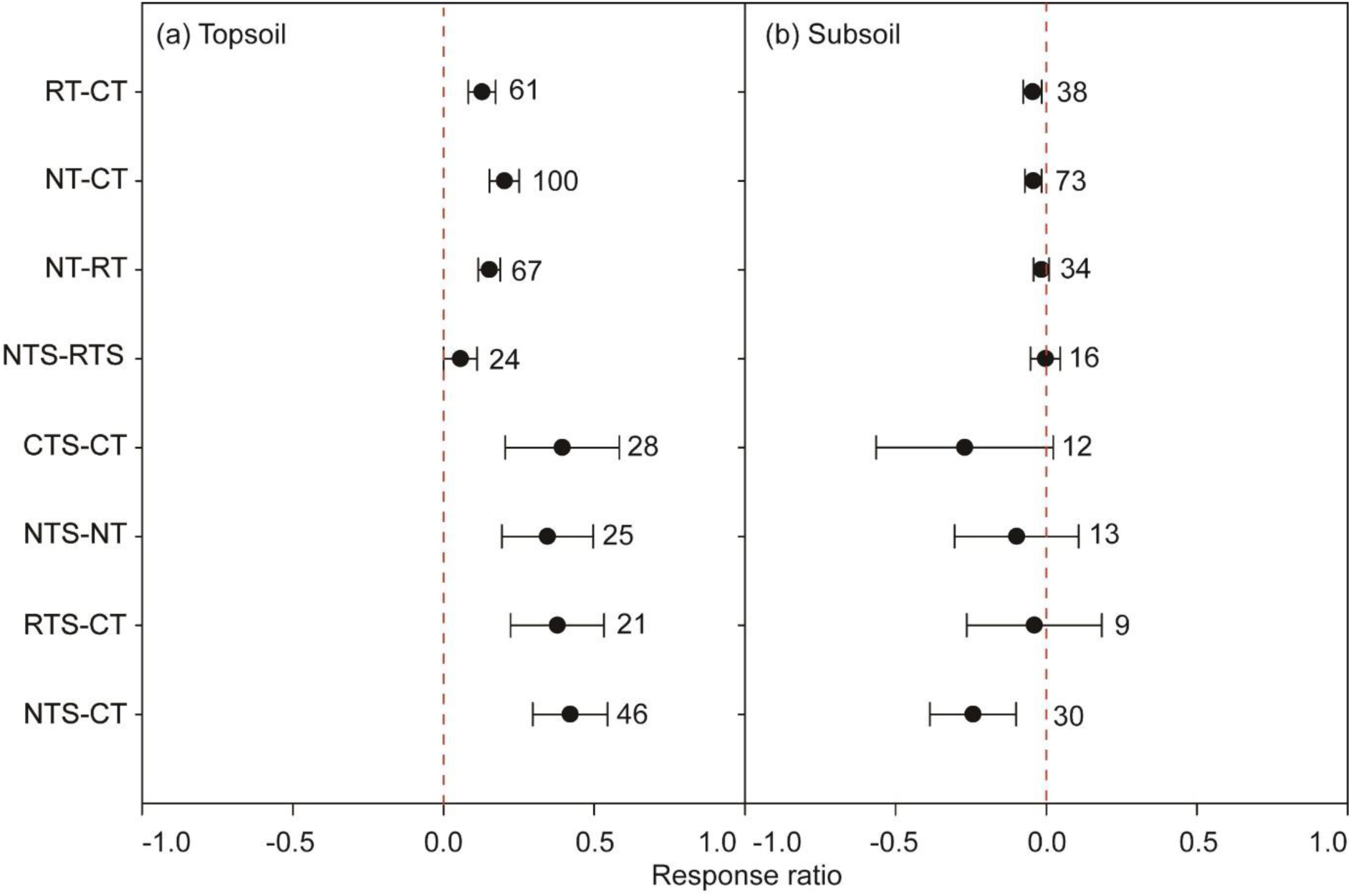
The effect size for soil organic carbon stock under different conservation agricultural practices at different soil sampling depths, a) topsoil, and b) subsoil. Effect size stands for the weighted response ratio between treatment and control. Error bars represent the 95% confidence intervals. The sample size of each variable is noted adjacent to each bar. For a description of the abbreviations, refer to the Fig. 2 title.

For sandy soils, NT significantly increased SOC stock as compared to CT (Fig. S1a), and NTS also significantly increased SOC stock as compared to CT. Whatever the conservation tillage practices, no significant increase was detected in loamy soils (Fig. S1b). For silty soils, RT, NT, RTS, and CTS significantly increased SOC stock relative to CT, but NTS significantly decreased SOC stock as compared to NT and CT, respectively (Fig. S1c). Only NT had significant effects on increasing SOC stock as compared to CT in clayey soils (Fig. S1d).

In semi-humid areas, no significant changes in SOC stocks were found for all comparisons (Fig. 6a). In most cases of humid areas, treatments increased SOC stock (Fig. 6b, *p* < 0.01), with the exceptions of NT as compared with RT and CTS as compared with CT. The increase in SOC stock was highest (by 417%) under RTS compared to CT. For the perhumid areas, NTS significantly decreased SOC stock as compared to NT (Fig. 6c), while the rest of the conservation tillage practices increased SOC stock (*p* < 0.01).

**Figure 6.**
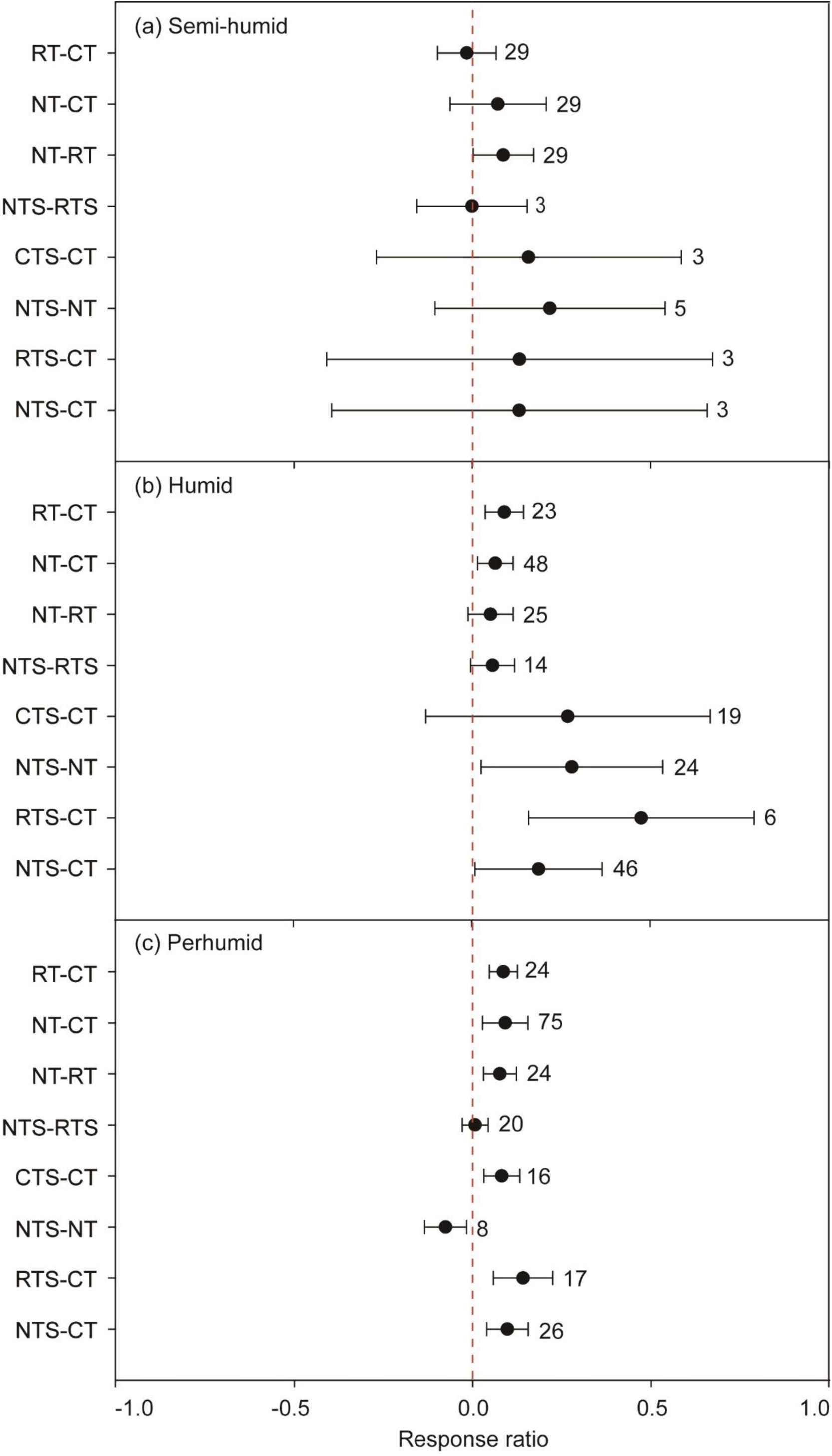
The effect size for soil organic carbon stock under different conservation agricultural practices across various climate categories as indicated by aridity index (AI), a) semi-humid, 10-20, b) humid, 20-30, c) perhumid, >30. Effect size stands for the weighted response ratio between treatment and control. Error bars represent the 95% confidence intervals. The sample size of each variable is noted adjacent to each bar. For a description of the abbreviations, refer to the Fig. 2 title.

### 3.4 Determining key drivers behind increased C stock under conservation tillage practices

The network graph reveals the interactions of C stock to soil properties, environmental and management conditions (Fig. 7). Initial SOC (coefficient = 0.39), TN (46) and MAP (0.29) had the highest positive correlation with the C stock. Additionally, cropping intensity (0.16), residue retention (0.16) and study duration (0.10) positively correlated with C stock. However, the initial soil available phosphorus content (AP, −0.17) and soil sandy content (−0.21) negatively correlated with C stock. Tillage practice affected the C stock (Fig. S1) while it did not show significant correlations with other variables and thus was excluded. Since initial SOC and TN were tightly correlated (r^2^ = 40, *p*<0.01), only the regression analysis between initial SOC and C stock was shown and C stock was positively correlated with initial SOC (r^2^ = 0.23, *p*<0.01, Fig. 8), C stock was also positively correlated with initial TN (r^2^ = 0.19, *p*<0.01), and MAP (r^2^ = 0.03, *p*=0.03, Fig. S2).

**Figure 7.**
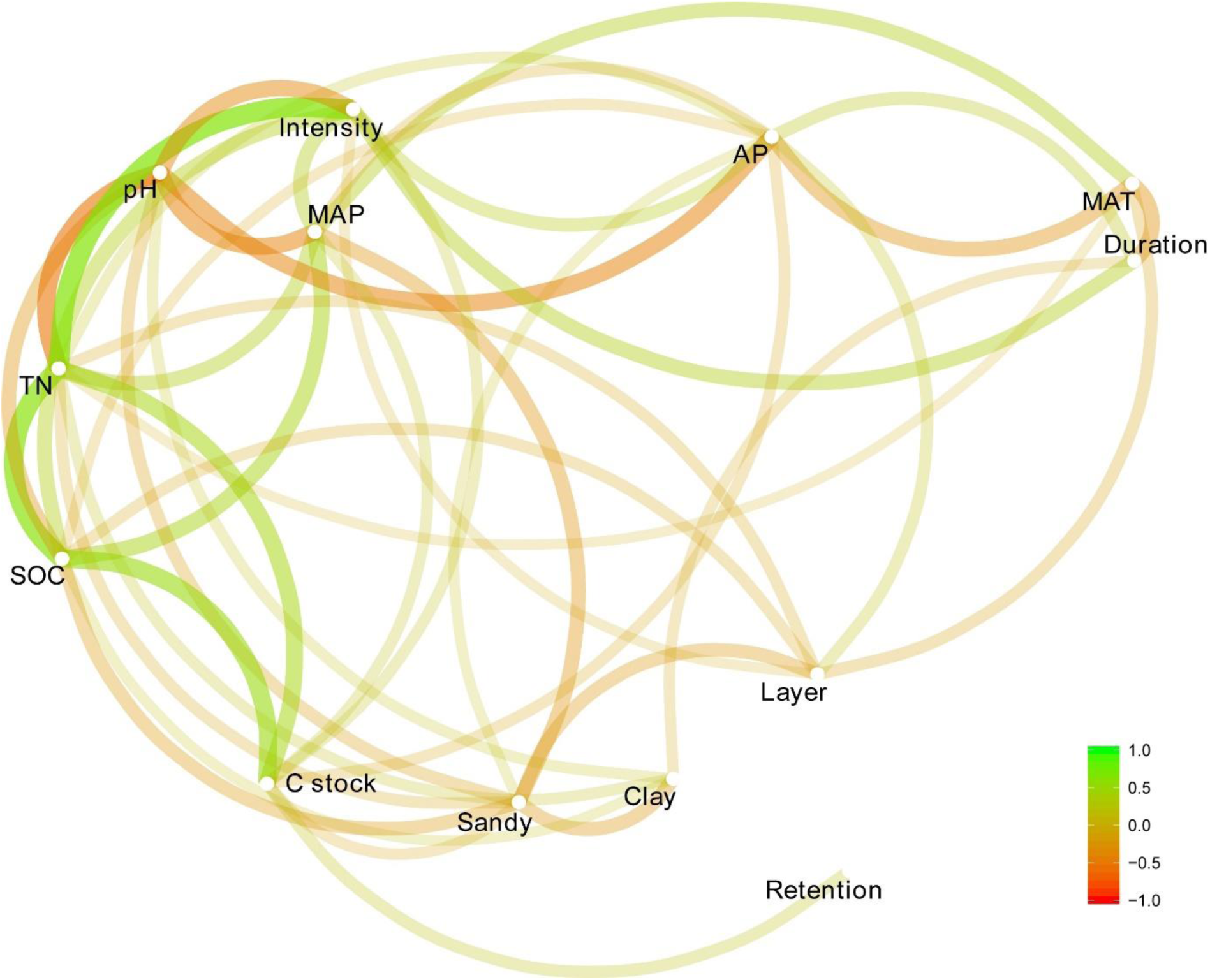
Empirical network of carbon (C) stock to soil properties, environmental and management conditions. A correlation coefficient is indicated by the colour gradient. AP, initial available phosphorus content; Clay, soil clay content; Duration, study duration; Intensity, cropping intensity; Layer, sampling depth; MAP, mean annual precipitation; MAT, mean annual temperature; Retention, residual retention; Sandy, soil sandy content; SOC, initial SOC content; TN, initial soil total nitrogen content (the colour version of the figure is in the web version of this article).

**Figure 8.**
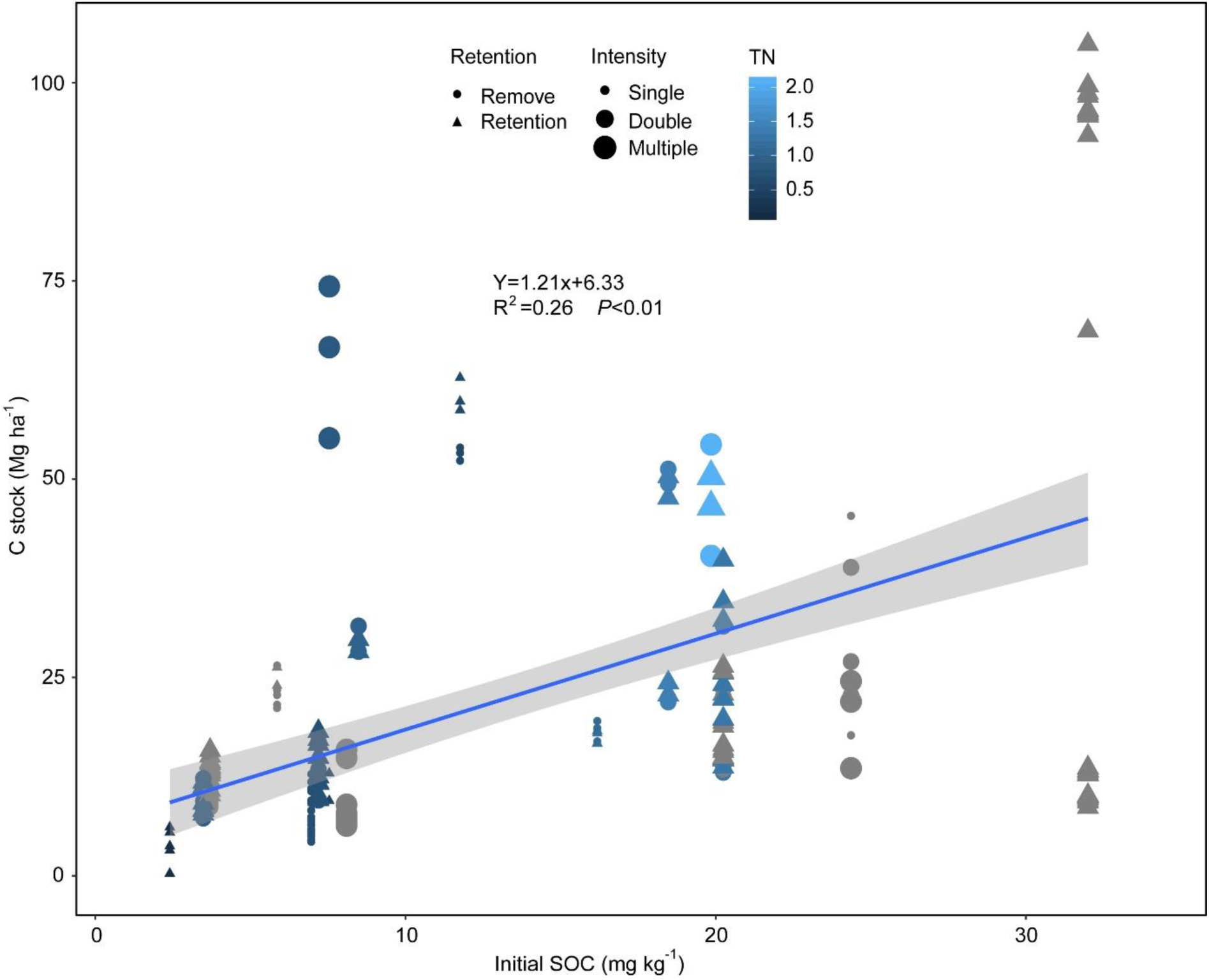
Relationships between soil initial soil organic carbon (SOC) and soil C stock. Initial soil total nitrogen content (TN) is indicated by the colour gradient. Management conditions, cropping intensity and residue retention are indicated by symbol shape and size, respectively (the colour version can be found in the web version of this article).

## 4. Discussion

### 4.1 Minimum tillage increases SOC stock

All minimum tillage practices that are considered in the present study show greater SOC stock in comparison with CT, which supports part of our hypothesis. The increased SOC is likely related to the redistribution of SOC to within aggregates which increases its stability under minimum tillage (Sheehy et al., 2015). The greater soil aggregation would slow organic matter decomposition rates in minimum tillage plots (Balkcom et al., 2013; Luo et al., 2017; Sauvadet et al., 2018).

The greater enhancement of SOC stock in the topsoil (0-15 cm) than in the subsoil (> 15 cm) is consistent with the positive effects of NT on SOC stock being mainly limited to surface soil layers reported in various studies (e.g., (Baker et al., 2007; Balkcom et al., 2013; Follett et al., 2013; Hubbard et al., 2013; Luo et al., 2010)). Salome et al. (2010) have shown that the physical accessibility of the organic C to microorganisms was a major control on C dynamics in the subsoil. The slow increase in SOC stock in the subsoil is thus probably due to the reduction of soil disturbance in NT and RT systems causing the vertical separation of decomposers and the substrate in the subsoil (Chenu et al., 2019; Li et al., 2018). Meanwhile, Salome et al. (2010) reported that spatial heterogeneity in C content was greater in the subsoil than in topsoil at the field scale. Collectively, subsoil has less potential in gaining SOC stock in the cropland.

In addition, NT increased more SOC stock in comparison with RT in the semi-humid regions. This finding is in line with a synthesis study in Spain, which indicated that the C sequestration rate under NT was 5 times higher than under RT (González-Sánchez et al., 2012). This is likely due to the capacity in water conservation under NT, and NT has been found to be more effective in reserving water in semi-humid areas (Busari et al., 2015), which facilitates root growth, nutrient use efficiencies and ultimately the agronomic yield (Pittelkow et al., 2014). Moreover, the C loss rates were also reported to be lower under NT than under RT (Fabrizzi et al., 2007), thereby resulted in higher SOC stock under NT than under RT.

Duration of minimum tillage practice also exhibited a strong influence on SOC stock. Our study found that NT increased SOC rapidly in the first 6 years, followed by a period of slower sequestration phase before reaching a slow steady rate. González-Sánchez et al. (2012) also found higher SOC sequestration rate during the initial years of conservation tillage practice based on 29 studies in Spain. In line with other case studies, the change of the C sequestration is related to the storage capacity of the soil associating to the duration of minimum tillage practices (Lu et al., 2009; Luo et al., 2010). In other words, the shorter the minimum tillage duration, the further a soil is from the upper limit, thereby the greater the potential for increasing SOC stocks.

In addition, more studies on soil N stock in responses to conservation tillage practices are needed to draw a more robust conclusion, where the 95% CI for N stock was very large due to insufficient data (Fig. 2b). Additionally, soil N is more variable across different cropping systems, and it is usually more sensitive than SOC to residue retention. Thus, the 95% CI for N stock was very large when residue retention was involved.

### 4.2 Minimum tillage coupled with residue retention is more effective in increasing SOC stock

Our analysis showed that residue retention facilitated the increase in SOC stock in the plots with minimum tillage. The potential mechanisms of residue return to increase SOC stock have been well documented. Firstly, in addition to increasing SOC by directly adding C into the soil, residue retention also improves soil physical and chemical properties (Johnson and Hoyt, 1999) and ultimately increases soil C storage (Pittelkow et al., 2014). Secondly, residue retention increases C stabilization by regulating microbial functions such as soil C- and N-cycling enzyme activities during litter degradation (Luo et al., 2018; Sauvadet et al., 2018; Zuber and Villamil, 2016), increases soil microbial biomass (Li et al., 2018), and enhances mineralization of nutrients (Cheng et al., 2017). Compared with minimum tillage practices alone, residue retention was more promising in increasing SOC stock (Han et al., 2018; Lu et al., 2009). Therefore, more organic C input and less frequent soil disturbance could lead to a greater SOC accumulation (Hubbard et al., 2013; Liu et al., 2014). However, it should be noted that our meta-analysis could not differentiate the method/form and rate of residue retention, which could also affect the direction and magnitude of the effect of residue retention on SOC stock (Han et al., 2018).

The combined effect of residue retention and minimum tillage was more obvious in the topsoil. Significant priming effect was observed in the topsoil (Salome et al., 2010), which indicated that the availability of labile C was the most important limiting factor on C dynamics in the topsoil. Residue retention alleviates the C limitation (Liu et al., 2014), and thus increases soil microbial population size (Li et al., 2018), which may enhance aggregate formation and stability (Novelli et al., 2017; Sheehy et al., 2015). High SOC stock naturally occurs in the topsoil under conservation tillage practices where there is more C input through residue retention and less output due to aggregation protection. Greater SOC accumulation in the topsoil from residue retention, and a negative relationship between the soil depth and the lnRR of SOC with straw return have been reported in a global synthesis (Liu et al. (2014). Our study further showed that residue retention increased SOC rapidly in the first 6 years. Furthermore, as compared to CT, incorporation of residue with minimum tillage consistently increased SOC stock when the study duration was less than 12 years, which is partly supported by the earlier finding that 12 years of minimum tillage resulted in C equilibrium in these systems (Lu et al., 2009; Luo et al., 2017).

Cropping intensity, on the other hand, alters the quality and quantity of residue-C input into the soil, thus, directly impacts SOC turnover (Liu et al., 2014; Novelli et al., 2017) and changes organic C forms that could support increased soil microbial biomass (Li et al., 2018; Somenahally et al., 2018) and soil fertility (Luo et al., 2018; Schmidt et al., 2018). Double cropping systems increased SOC stock in our study as compared to single cropping systems. However, the positive effect of minimum tillage or residue retention on SOC stock declined under multiple cropping intensity (> 3 crops a year), which could be due to the excessive soil disturbance during the planting operations and competition for water and nutrients (Novelli et al., 2017), and thereby increased C output through decomposition but decreased C input by lowering crop yield. Therefore, it can be concluded that only the increased retention of residue plus a reasonable increase in cropping intensity can increase soil C storage.

### 4.3 Soil properties were more critical than climatic condition and management practices in changing SOC stock

We found that climate conditions and management practices were not as important as initial soil nutrient availabilities in changing SOC stock (Fig. 7), consistent with some earlier studies (Smith et al., 2005). However, our result is in contrast with a South Africa long-term field trial, where seasonal weather variation was the main determinant of the impact of conservation tillage practices on SOC, irrespective of management and soil nutrient availabilities (Swanepoel et al., 2018). Overall, conservation tillage practices did not increase SOC stock in semi-humid climate conditions in our meta-analysis. As expected, SOC accumulated faster in the wetter zones than in the drier regions (Zhang et al., 2015). Increase in SOC stocks in Southern Africa with conservation tillage practices were lower than anticipated as a result of limited biomass production (Cheesman et al., 2016). The decomposition rate of SOC constants increased exponentially with the increase of MAT (Zhang et al., 2015), however, plant productivity was also likely to increase, which probably offset soil C losses. In most cases, conservation tillage practices increased SOC stock in humid or perhumid climate conditions. Similarly, crop growth and the consequent C input to soil tend to be higher in warmer or wetter regions than in colder or drier regions. However, soil microbial activities also increase in warmer or wetter regions (Wu et al., 2011; Zhang et al., 2015), leading to little SOC increase under all conservation tillage practices (Fig. 6b and 7c). In semi-arid regions, SOC stocks were usually less variable in the short-term as it requires several years to detect significant changes in microbial activities or SOM decomposition rates (Novelli et al., 2017). Ecosystem responses to the combination of temperature and altered precipitation tended to be smaller than expected from the single-factor responses (Wu et al., 2011). Part of the reason for this discrepancy was caused by the grouping/clustering of results based on AI (an index derived from MAT and MAP) in our analysis, instead of using soil temperature and water content at the time of sampling, which was mostly not reported in the publications used in this meta-analysis.

The merits of conservation tillage practice on SOC accumulation depends on soil conditions, especially the initial soil nutrient level. Initial soil nutrient conditions regulate the quality and quantity of plant C inputs to the soil by regulating aboveground biomass (Lal, 2004a; Luo et al., 2018). Soil microbe is critical for turnover and stabilization of soil organic matter (Salome et al., 2010; Sheehy et al., 2015), and the initial soil nutrient conditions play essential roles in changing the microbial community composition and C use efficiency (Sauvadet et al., 2018). In this study, the initial SOC stands out of other soil nutrient conditions, this partly because the important role of initial SOC concentration for regulating soil C formation and stability through direct and indirect effects on the turnover of aboveground litter, root C inputs, aggregate stability, and microbial process (Xu et al., 2018). Additionally, residue retention contributes more to SOC sequestration in coarse-textured than fine-textured soils (Wan et al., 2018), which was supported by our finding (Fig. S1a). Meanwhile, a 6 and 8 year conservation tillage field trials in South Africa found that sandy soils were more influenced by the climate than clayey soils on SOC changes (Swanepoel et al., 2018). Also, decreased tillage intensity increased SOC stock in silty or clayey soils (Fig. S1c, d). The finer soil particles play a vital role in increasing the cation exchange and water holding capacity, which are critical for C turnover by providing water and nutrients for microbial growth (Liu et al., 2014; Wan et al., 2018). Clay content has been considered a critical soil property to influence its capacity to store C (Novelli et al., 2017; Wan et al., 2018), and clayey soils often have higher initial SOC and also have greater capacity in increasing SOC stock.

## 5. Conclusions

Increasing SOC stock is critical for improving soil health, crop productivity, and agroecological resilience, as well as for mitigating greenhouse gas emission (Chenu et al., 2019). The conservation tillage practice has been acknowledged as a measure with a high potential for promoting SOC pool, it is thus important to assess the effect of different conservation tillage practices in enhancing the SOC reserve. Based on a global dataset, the meta-analysis showed that conservation tillage practices could generally increase SOC stock. Specifically, a combination of minimum tillage and residue addition is more effective than residue addition alone in increasing SOC stock, in turn, have higher efficiency than minimum tillage alone. All these effects were more obvious in the topsoil. However, conservation tillage practices are not effective in increasing SOC stock in arid, semi-arid, and semi-humid areas, thereby alternative management practices are warranted.

Moreover, initial soil nutrient availability and cropping intensity were found to be more critical than climatic conditions in influencing the effectiveness of conservation tillage on SOC stock. Double cropping generally increases SOC stock cross all conservation tillage practices as compared to single or multiple cropping. The positive initial SOC effect on cropland C stock highlights the requirement of soil C sequestration in order to enhance the potential in increasing C stock for coming generations. Both residue retention and NT increase SOC rapidly in the first 6 years regardless of soil texture or the climate region. The underlying mechanisms causing the variation in responses of SOC towards different external factors within different cropping systems need to be investigated using more detailed dataset. These results have important implications for sustainable soil management. We recommend that a combination of residue retention and minimum tillage in a double cropping agricultural system should be an important strategy to increase SOC storage in global farmlands.

## Acknowledgements

We thank all the researchers whose data were used in this meta-analysis. This work was jointly supported by the National Natural Science Foundation of China (31802133) and the China Scholarship Council (201606180104). The authors are very grateful for revisions to the manuscript by Huadong Zang (University of Göttingen).

## References

Adams, D.C., Gurevitch, J., Rosenberg, M.S., 1997. Resampling tests for meta-analysis of ecological data. Ecology 78, 1277–1283.

Baker, J.M., Ochsner, T.E., Venterea, R.T., Griffis, T.J., 2007. Tillage and soil carbon sequestration-What do we really know? Agric Ecosyst Environ. 118, 1–5.

Balkcom, K.S., Arriaga, F.J., van Santen, E., 2013. Conservation systems to enhance soil carbon sequestration in the Southeast US Coastal Plain. Soil Sci. Soc. Am. J. 77, 1774–1783.

Blanco-Canqui, H., Ruis, S.J., 2018. No-tillage and soil physical environment. Geoderma 326, 164–200.

Busari, M.A., Kukal, S.S., Kaur, A., Bhatt, R., Dulazi, A.A., 2015. Conservation tillage impacts on soil, crop and the environment. J. Soil Water Conserv. 3, 119–129.

Cheesman, S., Thierfelder, C., Eash, N.S., Kassie, G.T., Frossard, E., 2016. Soil carbon stocks in conservation agriculture systems of Southern Africa. Soi. Tillage. Res. 156, 99–109.

Cheng, Y., Wang, J., Wang, J., Chang, S.X., Wang, S., 2017. The quality and quantity of exogenous organic carbon input control microbial NO_3_^−^ immobilization: A meta-analysis. Soil Biol Biochem. 115, 357–363.

Chenu, C., Angers, D.A., Barré, P., Derrien, D., Arrouays, D., Balesdent, J., 2019. Increasing organic stocks in agricultural soils: Knowledge gaps and potential innovations. Soi. Tillage. Res. 188, 41–52.

De Martonne, E., 1926. Regions of Interior Basin Drainage. Geogr. Rev. 182, 1395–1398.

Fabrizzi, K., Rice, C.W., Izaurralde, R.C., 2007. Soil carbon sequestration in Kansas: Long-term effect of tillage, N fertilization, and crop rotation, The Fourth USDA Greenhouse Gas Conference, Marriott Hotel, Stadium Ballroom, Second Floor.

FAO, 2015. Permanent cropland, World Bank Open Data.

Follett, R.F., Jantalia, C.P., Halvorson, A.D., 2013. Soil carbon dynamics for irrigated corn under two tillage systems. Soil Sci. Soc. Am. J. 77, 951–963.

González-Sánchez, E.J., Ordóñez-Fernández, R., Carbonell-Bojollo, R., Veroz-González, O., Gil-Ribes, J.A., 2012. Meta-analysis on atmospheric carbon capture in Spain through the use of conservation agriculture. Soi. Tillage. Res. 122, 52–60.

Gurevitch, J., Hedges, L., 2001. Meta-analysis: combining the results of independent experiments., In: Scheiner, S., Gurevitch, J. (Eds.), Design and analysis of ecological experiments. Oxford University Press, pp. 347–369.

Han, X., Xu, C., Dungait, J.A.J., Bol, R., Wang, X.J., Wu, W.L., Meng, F.Q., 2018. Straw incorporation increases crop yield and soil organic carbon sequestration but varies under different natural conditions and farming practices in China: a system analysis. Biogeosciences 15, 1933–1946.

Hedges, L.V., Gurevitch, J., Curtis, P.S., 1999. The meta-analysis of response ratios in experimental ecology. Ecology 80, 1150–1156.

Hubbard, R.K., Strickland, T.C., Phatak, S., 2013. Effects of cover crop systems on soil physical properties and carbon/nitrogen relationships in the coastal plain of southeastern USA. Soi. Tillage. Res. 126, 276–283.

Jackson, S., 2016. corrr: Correlations in R. R package version 0.2. 1. R Project.

Johnson, A.M., Hoyt, G.D., 1999. Changes to the soil environment under conservation tillage. HortTechnology 9, 380–393.

Lal, R., 2004a. Soil carbon sequestration impacts on global climate change and food security. Science 304, 1623–1627.

Lal, R., 2004b. Soil carbon sequestration to mitigate climate change. Geoderma 123, 1–22.

Li, Y., Chang, S.X., Tian, L.H., Zhang, Q.P., 2018. Conservation agriculture practices increase soil microbial biomass carbon and nitrogen in agricultural soils: A global meta-analysis. Soil Biol Biochem. 121, 50–58.

Liu, C., Lu, M., Cui, J., Li, B., Fang, C., 2014. Effects of straw carbon input on carbon dynamics in agricultural soils: a meta-analysis. Glob Chang Biol. 20, 1366–1381.

Lu, F., Wang, X., Han, B., Ouyang, Z., Duan, X., Zheng, H., Miao, H., 2009. Soil carbon sequestrations by nitrogen fertilizer application, straw return and no-tillage in China’s cropland. Glob Chang Biol. 15, 281–305.

Luo, G., Li, L., Friman, V.-P., Guo, J., Guo, S., Shen, Q., Ling, N., 2018. Organic amendments increase crop yields by improving microbe-mediated soil functioning of agroecosystems: A meta-analysis. Soil Biol Biochem. 124, 105–115.

Luo, Y., Hui, D., Zhang, D., 2006. Elevated CO2 stimulates net accumulations of carbon and nitrogen in land ecosystems: a meta-analysis. Ecology 87, 53–63.

Luo, Z., Feng, W., Luo, Y., Baldock, J., Wang, E., 2017. Soil organic carbon dynamics jointly controlled by climate, carbon inputs, soil properties and soil carbon fractions. Glob Chang Biol. 23, 4430–4439.

Luo, Z., Wang, E., Sun, O.J., 2010. Can no-tillage stimulate carbon sequestration in agricultural soils? A meta-analysis of paired experiments. Agric Ecosyst Environ. 139, 224–231.

Novelli, L.E., Caviglia, O.P., Piñeiro, G., 2017. Increased cropping intensity improves crop residue inputs to the soil and aggregate-associated soil organic carbon stocks. Soi. Tillage. Res. 165, 128–136.

Osenberg, C.W., Sarnelle, O., Cooper, S.D., Holt, R.D., 1999. Resolving ecological questions through meta-analysis: goals, metrics, and models. Ecology 80, 1105–1117.

Pittelkow, C.M., Liang, X., Linquist, B.A., van Groenigen, K.J., Lee, J., Lundy, M.E., van Gestel, N., Six, J., Venterea, R.T., van Kessel, C., 2014. Productivity limits and potentials of the principles of conservation agriculture. Nature 517, 365.

Poeplau, C., Don, A., 2015. Carbon sequestration in agricultural soils via cultivation of cover crops-A meta-analysis. Agric Ecosyst Environ. 200, 33–41.

Salome, C., Nunan, N., Pouteau, V., Lerch, T.Z., Chenu, C., 2010. Carbon dynamics in topsoil and in subsoil may be controlled by different regulatory mechanisms. Glob Chang Biol. 16, 416–426.

Sauvadet, M., Lashermes, G., Alavoine, G., Recous, S., Chauvat, M., Maron, P.A., Bertrand, I., 2018. High carbon use efficiency and low priming effect promote soil C stabilization under reduced tillage. Soil Biol Biochem. 123, 64–73.

Schmidt, E.S., Villamil, M.B., Amiotti, N.M., 2018. Soil quality under conservation practices on farm operations of the southern semiarid pampas region of Argentina. Soi. Tillage. Res. 176, 85–94.

Sheehy, J., Regina, K., Alakukku, L., Six, J., 2015. Impact of no-till and reduced tillage on aggregation and aggregate-associated carbon in Northern European agroecosystems. Soi. Tillage. Res. 150, 107–113.

Smith, P., Andrén, O., Karlsson, T., Perälä, P., Regina, K., Rounsevell, M., Wesemael, B., 2005. Carbon sequestration potential in European croplands has been overestimated. Glob Chang Biol. 11, 2153–2163.

Somenahally, A., DuPont, J.I., Brady, J., McLawrence, J., Northup, B., Gowda, P., 2018. Microbial communities in soil profile are more responsive to legacy effects of wheat-cover crop rotations than tillage systems. Soil Biol Biochem. 123, 126–135.

Strassmann, K.M., Joos, F., Fischer, G., 2008. Simulating effects of land use changes on carbon fluxes: past contributions to atmospheric CO2 increases and future commitments due to losses of terrestrial sink capacity. Tellus B 60, 583–603.

Swanepoel, C.M., Rötter, R.P., van der Laan, M., Annandale, J.G., Beukes, D.J., du Preez, C.C., Swanepoel, L.H., van der Merwe, A., Hoffmann, M.P., 2018. The benefits of conservation agriculture on soil organic carbon and yield in southern Africa are site-specific. Soi. Tillage. Res. 183, 72–82.

VandenBygaart, A.J., Gregorich, E.G., Angers, D.A., 2003. Influence of agricultural management on soil organic carbon: A compendium and assessment of Canadian studies. Can. J. Soil Sci. 83, 363–380.

Viechtbauer, W., 2010. Conducting meta-analyses in R with the metafor package. J Stat Softw. 36, 1–48.

Wall, P., 2006. Facilitating the widespread adoption of conservation agriculture and other resource conserving technologies (RCT’s): Some diffi cult issues. Science Week Extended Abstract. CIMMYT Headquarters, El Batan, Mexico, 61–64.

Wan, X., Xiao, L., Vadeboncoeur, M.A., Johnson, C.E., Huang, Z., 2018. Response of mineral soil carbon storage to harvest residue retention depends on soil texture: A meta-analysis. Forest Ecol Manag. 408, 9–15.

Wickham, H., 2016. ggplot2: elegant graphics for data analysis. Springer, New York.

Wu, Z., Dijkstra, P., George, W.K., Peñuelas, J., Hungate, B. A.,, 2011. Responses of terrestrial ecosystems to temperature and precipitation change: a meta-analysis of experimental manipulation. Glob Chang Biol. 17, 927–942.

Xu, S., Li, P., Sayer, E.J., Zhang, B., Wang, J., Qiao, C., Peng, Z., Diao, L., Chi, Y., Liu, W., Liu, L., 2018. Initial soil organic matter content influences the storage and turnover of litter, root and soil carbon in grasslands. Ecosystems 21, 1377–1389.

Zhang, K., Dang, H., Zhang, Q., Cheng, X., 2015. Soil carbon dynamics following land-use change varied with temperature and precipitation gradients: evidence from stable isotopes. Glob Chang Biol. 21, 2762–2772.

Zuber, S.M., Villamil, M.B., 2016. Meta-analysis approach to assess effect of tillage on microbial biomass and enzyme activities. Soil Biol Biochem. 97, 176–187.

