## Supplemental materials for "Residue retention promotes soil carbon accumulation in minimum tillage systems: Implications for conservation tillage"

**Table S1.** Between-group heterogeneity (Qb) and probability (P) among n observations of nitrogen stock in response to different conservation agriculture practices under different sampling depth. For a description of the abbreviations, refer Fig. 2 title.

|  | Test of heterogeneity | | | Random effects model | |
| --- | --- | --- | --- | --- | --- |
|  | Q | n | *p*-value | *z* | *p*-value |
| PT-RT | 336 | 40 | < .0001 | 2.6727 | 0.0075 |
| PT-NT | 796 | 86 | < .0001 | 4.2954 | <.0001 |
| PT-RTS | 1372 | 16 | < .0001 | 0.5140 | 0.6072 |
| PT-NTS | 4515 | 46 | < .0001 | 1.8621 | 0.0626 |
| PT-PTS | 3369 | 22 | < .0001 | 0.8721 | 0.3832 |
| RT-NT | 157 | 42 | < .0001 | 4.2547 | <.0001 |
| RT-RTS | 521 | 8 | < .0001 | -0.1715 | 0.8638 |
| RT-NTS | 545 | 7 | < .0001 | 0.5033 | 0.6148 |
| NT-NTS | 1960 | 19 | < .0001 | 2.2001 | 0.0278 |
| NT-RTS | 144 | 6 | < .0001 | 1.5754 | 0.1152 |
| PTS-RTS | 36 | 23 | 0.0279 | 1.5894 | 0.1120 |
| PTS-NTS | 1027 | 48 | 0.2055 | 1.2661 | 0.2055 |
| PTS-RT | 566 | 8 | < .0001 | -0.4048 | 0.6856 |
| PTS-NT | 1888 | 13 | < .0001 | -1.0532 | 0.2923 |
| RTS-NTS | 112 | 21 | < .0001 | 3.1411 | 0.0017 |

**Table S2.** Between-group heterogeneity (Qb) and probability (P) among n observations of carbon stock under different conservation agriculture practices. For a description of the abbreviations, refer Fig. 2 title.

| CAP | Test of heterogeneity | | | Random effects model | |
| --- | --- | --- | --- | --- | --- |
|  | Q | n | *p*-value | *z* | *p*-value |
| PT-RT | 1204 | 99 | < .0001 | 4.8878 | <.0001 |
| PT-NT | 62534 | 177 | < .0001 | 4.6783 | <.0001 |
| PT-RTS | 2756 | 30 | < .0001 | 5.4786 | <.0001 |
| PT-NTS | 5954 | 76 | < .0001 | 5.4426 | <.0001 |
| PT-PTS | 3994 | 40 | <.0001 | 4.8210 | <.0001 |
| RT-NT | 2065 | 101 | < .0001 | 7.4050 | <.0001 |
| RT-RTS | 2206 | 20 | <.0001 | 4.7743 | <.0001 |
| RT-NTS | 1118 | 10 | <.0001 | 1.0787 | 0.2807 |
| NT-NTS | 5608 | 38 | <.0001 | 6.3632 | <.0001 |
| NT-RTS | 720 | 12 | < .0001 | 2.0058 | 0.0449 |
| PTS-RTS | 384 | 48 | < .0001 | 2.7116 | 0.0067 |
| PTS-NTS | 2885 | 102 | < .0001 | 5.2339 | <.0001 |
| PTS-RT | 1121 | 8 | <.0001 | -2.6161 | 0.0089 |
| PTS-NT | 3040 | 22 | <.0001 | -5.8086 | <.0001 |
| RTS-NTS | 1820 | 40 | < .0001 | 1.7304 | 0.0836 |

**Table S3.** Between-group heterogeneity (Qb) and probability (P) among n observations of carbon stock in response to different conservation agriculture practices under various soil textures. For a description of the abbreviations, refer Fig. 2 title.

| PT-RT | Test of heterogeneity | | | Random effects model | |
| --- | --- | --- | --- | --- | --- |
|  | Q | n | *p*-value | *z* | *p*-value |
| Sandy | 474 | 26 | < .0001 | 0.5721 | 0.5672 |
| Silty | 441 | 25 | < .0001 | 2.4070 | 0.0161 |
| Loamy | 60 | 23 | < .0001 | 1.5199 | 0.1285 |
| Clayey | 70 | 19 | < .0001 | 5.4265 | <.0001 |

| PT-NT | Test of heterogeneity | | | Random effects model | |
| --- | --- | --- | --- | --- | --- |
|  | Q | n | *p*-value | *z* | *p*-value |
| Sandy | 510 | 46 | < .0001 | 2.1596 | 0.0308 |
| Silty | 3650 | 50 | <.0001 | 4.6221 | <.0001 |
| Loamy | 23851 | 52 | < .0001 | 1.7030 | 0.0886 |
| Clayey | 526 | 25 | < .0001 | 5.2505 | <.0001 |

| RT-NT | Test of heterogeneity | | | Random effects model | |
| --- | --- | --- | --- | --- | --- |
|  | Q | n | *p*-value | *z* | *p*-value |
| Sandy | 224 | 32 | < .0001 | 2.7164 | 0.0066 |
| Silty | 1482 | 26 | <.0001 | 6.7929 | <.0001 |
| Loamy | 75 | 25 | < .0001 | 1.2056 | 0.2280 |
| Clayey | 236 | 15 | < .0001 | 1.7690 | 0.0769 |

| PTS-RTS | Test of heterogeneity | | | Random effects model | |
| --- | --- | --- | --- | --- | --- |
|  | Q | n | *p*-value | *z* | *p*-value |
| Sandy | 42 | 11 | < .0001 | -1.0181 | 0.3086 |
| Silty | 185 | 6 | < .0001 | -0.4597 | 0.6457 |
| Loamy | 77 | 15 | < .0001 | 2.2941 | 0.0218 |
| Clayey | 41 | 13 | < .0001 | 1.8157 | 0.0694 |

| PTS-NTS | Test of heterogeneity | | | Random effects model | |
| --- | --- | --- | --- | --- | --- |
|  | Q | n | *p*-value | *z* | *p*-value |
| Sandy | 13 | 17 | 0.6644 | 0.1796 | 0.8575 |
| Silty | 515 | 27 | < .0001 | 5.6137 | < .0001 |
| Loamy | 1148 | 39 | < .0001 | 1.0896 | 0.2759 |
| Clayey | 67 | 14 | < .0001 | 2.1399 | 0.0324 |

| RTS-NTS | Test of heterogeneity | | | Random effects model | |
| --- | --- | --- | --- | --- | --- |
|  | Q | n | *p*-value | *z* | *p*-value |
| Sandy | 2 | 4 | 0.6890 | 4.9997 | <.0001 |
| Silty | 42 | 6 | <.0001 | -1.6306 | 0.1030 |
| Loamy | 80 | 13 | < .0001 | 1.8478 | 0.0646 |
| Clayey | 354 | 14 | < .0001 | -0.0575 | 0.9541 |

| PT-RTS | Test of heterogeneity | | | Random effects model | |
| --- | --- | --- | --- | --- | --- |
|  | Q | n | *p*-value | *z* | *p*-value |
| Sandy | 806 | 5 | < .0001 | 0.9244 | 0.3553 |
| Silty | 229 | 6 | <.0001 | 2.1792 | 0.0293 |
| Loamy | 102 | 8 | < .0001 | 1.7869 | 0.0739 |
| Clayey | 1163 | 8 | < .0001 | 1.9253 | 0.0542 |

| PT-NTS | Test of heterogeneity | | | Random effects model | |
| --- | --- | --- | --- | --- | --- |
|  | Q | n | *p*-value | *z* | *p*-value |
| Sandy | 2822 | 26 | < .0001 | 2.0231 | 0.0431 |
| Silty | 887 | 12 | <.0001 | -2.7697 | 0.0056 |
| Loamy | 786 | 17 | < .0001 | 1.0805 | 0.2799 |
| Clayey | 666 | 18 | < .0001 | 1.1417 | 0.2536 |

| PT-PTS | Test of heterogeneity | | | Random effects model | |
| --- | --- | --- | --- | --- | --- |
|  | Q | n | *p*-value | *z* | *p*-value |
| Sandy | 1958 | 16 | < .0001 | 1.9472 | 0.0515 |
| Silty | 49 | 5 | <.0001 | 3.0516 | 0.0023 |
| Loamy | 104 | 5 | < .0001 | -0.1877 | 0.8511 |
| Clayey | 1185 | 11 | < .0001 | 0.1485 | 0.8819 |

| NT-NTS | Test of heterogeneity | | | Random effects model | |
| --- | --- | --- | --- | --- | --- |
|  | Q | n | *p*-value | *z* | *p*-value |
| Sandy | 1807 | 16 | < .0001 | 1.8248 | 0.0680 |
| Silty | 477 | 3 | <.0001 | -6.6399 | <.0001 |
| Loamy | 1306 | 10 | < .0001 | 1.9110 | 0.0560 |
| Clayey | 474 | 8 | < .0001 | 0.4090 | 0.6825 |

**Table S4.** Between-group heterogeneity (Qb) and probability (P) among n observations of carbon stock in response to different conservation agriculture practices under various study duration. For a description of the abbreviations, refer Fig. 2 title.

| PT-RT | Test of heterogeneity | | | Random effects model | |
| --- | --- | --- | --- | --- | --- |
|  | Q | n | *p*-value | *z* | *p*-value |
| < 6 | 352 | 25 | < .0001 | -1.2641 | 0.2062 |
| 6 – 12 | 301 | 25 | < .0001 | 3.5651 | 0.0004 |
| > 12 | 365 | 43 | < .0001 | 2.1900 | 0.0285 |

| PT-NT | Test of heterogeneity | | | Random effects model | |
| --- | --- | --- | --- | --- | --- |
|  | Q | n | *p*-value | *z* | *p*-value |
| < 6 | 1122 | 42 | < .0001 | 7.4547 | <.0001 |
| 6 – 12 | 27074 | 52 | < .0001 | 1.8753 | 0.0607 |
| > 12 | 3980 | 79 | < .0001 | 3.5566 | 0.0004 |

| RT-NT | Test of heterogeneity | | | Random effects model | |
| --- | --- | --- | --- | --- | --- |
|  | Q | n | *p*-value | *z* | *p*-value |
| < 6 | 1246 | 24 | < .0001 | 6.7743 | <.0001 |
| 6 – 12 | 243 | 18 | < .0001 | 1.7620 | 0.0781 |
| > 12 | 530 | 59 | < .0001 | 2.5199 | 0.0117 |

| PTS-RTS | Test of heterogeneity | | | Random effects model | |
| --- | --- | --- | --- | --- | --- |
|  | Q | n | *p*-value | *z* | *p*-value |
| < 6 | 290 | 17 | < .0001 | 1.9083 | 0.0564 |
| 6 – 12 | 77 | 23 | < .0001 | 0.9317 | 0.3515 |
| > 12 | 12 | 5 | 0.0154 | 1.3672 | 0.1716 |

| PTS-NTS | Test of heterogeneity | | | Random effects model | |
| --- | --- | --- | --- | --- | --- |
|  | Q | n | *p*-value | *z* | *p*-value |
| < 6 | 475 | 27 | < .0001 | 4.2157 | <.0001 |
| 6 – 12 | 1207 | 56 | < .0001 | 1.1119 | 0.2662 |
| > 12 | 74 | 14 | < .0001 | 3.3937 | 0.0007 |

| RTS-NTS | Test of heterogeneity | | | Random effects model | |
| --- | --- | --- | --- | --- | --- |
|  | Q | n | *p*-value | *z* | *p*-value |
| < 6 | 196 | 16 | < .0001 | 1.2975 | 0.1944 |
| 6 – 12 | 183 | 21 | < .0001 | 0.425 | 0.6708 |
| > 12 | 5 | 3 | 0.0833 | 2.9806 | 0.0029 |

| PT-RTS | Test of heterogeneity | | | Random effects model | |
| --- | --- | --- | --- | --- | --- |
|  | Q | n | *p*-value | *z* | *p*-value |
| < 6 | 778 | 10 | < .0001 | 3.2223 | 0.0013 |
| 6 – 12 | 1474 | 16 | < .0001 | 2.6745 | 0.0075 |
| > 12 | 45 | 4 | < .0001 | 1.1942 | 0.2324 |

| PT-NTS | Test of heterogeneity | | | Random effects model | |
| --- | --- | --- | --- | --- | --- |
|  | Q | n | *p*-value | *z* | *p*-value |
| < 6 | 524 | 15 | < .0001 | 9.7602 | <.0001 |
| 6 – 12 | 4539 | 54 | < .0001 | 2.0382 | 0.0415 |
| > 12 | 521 | 7 | < .0001 | -0.8901 | 0.3734 |

| PT-PTS | Test of heterogeneity | | | Random effects model | |
| --- | --- | --- | --- | --- | --- |
|  | Q | n | *p*-value | *z* | *p*-value |
| < 6 | 98 | 7 | < .0001 | 4.9520 | <.0001 |
| 6 – 12 | 2618 | 29 | < .0001 | 2.3524 | 0.0187 |
| > 12 | 135 | 4 | < .0001 | -2.0355 | 0.0418 |

| NT-NTS | Test of heterogeneity | | | Random effects model | |
| --- | --- | --- | --- | --- | --- |
|  | Q | n | *p*-value | *z* | *p*-value |
| < 6 | 2139 | 12 | < .0001 | 5.9838 | <.0001 |
| 6 – 12 | 3192 | 24 | < .0001 | 1.8764 | 0.0606 |
| > 12 | 2 | 2 | 0.1716 | -12.44 | <.0001 |

**Table S5.** Between-group heterogeneity (Qb) and probability (P) among n observations of carbon stock in response to different conservation agriculture practices under various cropping intensity. For a description of the abbreviations, refer Fig. 2 title.

| PT-RT | Test of heterogeneity | | | Random effects model | |
| --- | --- | --- | --- | --- | --- |
|  | Q | n | *p*-value | *z* | *p*-value |
| Single | 390 | 32 | < .0001 | -1.0781 | 0.2810 |
| Double | 245 | 41 | < .0001 | 5.4607 | <.0001 |
| More | 368 | 20 | < .0001 | 0.9515 | 0.3413 |

| PT-NT | Test of heterogeneity | | | Random effects model | |
| --- | --- | --- | --- | --- | --- |
|  | Q | n | *p*-value | *z* | *p*-value |
| Single | 1210 | 42 | < .0001 | 5.5143 | <.0001 |
| Double | 5003 | 77 | < .0001 | 4.2180 | <.0001 |
| More | 20673 | 54 | < .0001 | 2.4737 | 0.0134 |

| RT-NT | Test of heterogeneity | | | Random effects model | |
| --- | --- | --- | --- | --- | --- |
|  | Q | n | *p*-value | *z* | *p*-value |
| Single | 1390 | 33 | < .0001 | 6.7345 | <.0001 |
| Double | 457 | 33 | < .0001 | 2.7474 | 0.0060 |
| More | 179 | 35 | < .0001 | 1.7867 | 0.0740 |

| PTS-RTS | Test of heterogeneity | | | Random effects model | |
| --- | --- | --- | --- | --- | --- |
|  | Q | n | *p*-value | *z* | *p*-value |
| Single | 206 | 8 | < .0001 | 0.2487 | 0.8036 |
| Double | 156 | 32 | < .0001 | 1.9093 | 0.0562 |
| More | 1 | 5 | 0.8497 | -0.1389 | 0.8895 |

| PTS-NTS | Test of heterogeneity | | | Random effects model | |
| --- | --- | --- | --- | --- | --- |
|  | Q | n | *p*-value | *z* | *p*-value |
| Single | 332 | 10 | < .0001 | 3.8876 | 0.0001 |
| Double | 1336 | 74 | < .0001 | 2.1389 | 0.0324 |
| More | 89 | 13 | < .0001 | -0.2264 | 0.8209 |

| RTS-NTS | Test of heterogeneity | | | Random effects model | |
| --- | --- | --- | --- | --- | --- |
|  | Q | n | *p*-value | *z* | *p*-value |
| Single | 121 | 5 | < .0001 | -0.0701 | 0.9441 |
| Double | 257 | 31 | < .0001 | 1.0281 | 0.3039 |
| More | 11 | 4 | 0.0117 | 1.5440 | 0.1226 |

| PT-RTS | Test of heterogeneity | | | Random effects model | |
| --- | --- | --- | --- | --- | --- |
|  | Q | n | *p*-value | *z* | *p*-value |
| Single | 247 | 5 | < .0001 | 4.5599 | <.0001 |
| Double | 2058 | 24 | < .0001 | 2.2360 | 0.0254 |
| More | 0 | 1 | 1 | 12.6680 | <.0001 |

| PT-NTS | Test of heterogeneity | | | Random effects model | |
| --- | --- | --- | --- | --- | --- |
|  | Q | n | *p*-value | *z* | *p*-value |
| Single | 2163 | 29 | < .0001 | 0.5833 | 0.5597 |
| Double | 2254 | 41 | < .0001 | 3.0178 | 0.0025 |
| More | 1134 | 6 | < .0001 | 0.942 | 0.3458 |
| PT-PTS | Test of heterogeneity | | | Random effects model | |
|  | Q | n | *p*-value | *z* | *p*-value |
| Single | 203 | 9 | < .0001 | -1.8658 | 0.0621 |
| Double | 1840 | 25 | < .0001 | 1.9883 | 0.0468 |
| More | 1176 | 6 | < .0001 | 0.9439 | 0.3452 |

| NT-NTS | Test of heterogeneity | | | Random effects model | |
| --- | --- | --- | --- | --- | --- |
|  | Q | n | *p*-value | *z* | *p*-value |
| Single | 482 | 5 | < .0001 | -5.8550 | <.0001 |
| Double | 1927 | 22 | < .0001 | 2.2775 | 0.0228 |
| More | 2175 | 11 | < .0001 | 2.0725 | 0.0382 |
| PT-RT | Test of heterogeneity | | | Random effects | |

**Table S6.** Between-group heterogeneity (Qb) and probability (P) among n observations of carbon stock in response to different conservation agriculture practices under various aridity index. For a description of the abbreviations, refer Fig. 2 title.

| PT-RT | Test of heterogeneity | | | Random effects model | |
| --- | --- | --- | --- | --- | --- |
|  | Q | n | *p*-value | *z* | *p*-value |
| 0-10 |  |  |  |  |  |
| 10-20 | 216 | 29 | < .0001 | 0.7220 | 0.7220 |
| 20-30 | 156 | 23 | < .0001 | 3.3529 | 0.0008 |
| >30 | 580 | 24 | < .0001 | 4.2978 | <.0001 |

| PT-NT | Test of heterogeneity | | | Random effects model | |
| --- | --- | --- | --- | --- | --- |
|  | Q | n | *p*-value | *z* | *p*-value |
| 0-10 |  |  |  |  |  |
| 10-20 | 575 | 29 | < .0001 | 1.0535 | 0.2921 |
| 20-30 | 588 | 48 | < .0001 | 2.6299 | 0.0085 |
| >30 | 60182 | 75 | < .0001 | 2.8203 | 0.0048 |

| RT-NT | Test of heterogeneity | | | Random effects model | |
| --- | --- | --- | --- | --- | --- |
|  | Q | n | *p*-value | *z* | *p*-value |
| 0-10 |  |  |  |  |  |
| 10-20 | 229 | 29 | < .0001 | 2.0149 | 0.0439 |
| 20-30 | 280 | 25 | < .0001 | 1.6522 | 0.0985 |
| >30 | 1160 | 24 | < .0001 | 3.2681 | 0.0011 |

| PTS-RTS | Test of heterogeneity | | | Random effects model | |
| --- | --- | --- | --- | --- | --- |
|  | Q | n | *p*-value | *z* | *p*-value |
| 0-10 | 0 | 1 | 1 | 0.7341 | 0.4629 |
| 10-20 | 6 | 4 | 0.1243 | -0.7775 | 0.4369 |
| 20-30 | 103 | 17 | 0.8497 | < .0001 | 0.7164 |
| >30 | 265 | 24 | < .0001 | 2.8167 | 0.0049 |

| PTS-NTS | Test of heterogeneity | | | Random effects model | |
| --- | --- | --- | --- | --- | --- |
|  | Q | n | *p*-value | *z* | *p*-value |
| 0-10 | 0 | 1 | 1 | 1.3293 | 0.1838 |
| 10-20 | 61 | 10 | < .0001 | -0.9220 | 0.3565 |
| 20-30 | 201 | 36 | < .0001 | 2.9348 | 0.0033 |
| >30 | 1577 | 54 | < .0001 | 5.3347 | <.0001 |

| RTS-NTS | Test of heterogeneity | | | Random effects model | |
| --- | --- | --- | --- | --- | --- |
|  | Q | n | *p*-value | *z* | *p*-value |
| 0-10 | 0 | 1 | 1 | 0.5948 | 0.5520 |
| 10-20 | 6 | 3 | 0.0608 | -0.0070 | 0.9944 |
| 20-30 | 354 | 14 | < .0001 | 1.8626 | 0.0625 |
| >30 | 151 | 20 | < .0001 | 0.3882 | 0.6979 |

| PT-RTS | Test of heterogeneity | | | Random effects model | |
| --- | --- | --- | --- | --- | --- |
|  | Q | n | *p*-value | *z* | *p*-value |
| 0-10 |  |  |  |  |  |
| 10-20 | 69 | 3 | < .0001 | 0.4815 | 0.6301 |
| 20-30 | 505 | 6 | < .0001 | 2.9521 | 0.0032 |
| >30 | 916 | 17 | < .0001 | 3.3469 | 0.0008 |

| PT-NTS | Test of heterogeneity | | | Random effects model | |
| --- | --- | --- | --- | --- | --- |
|  | Q | n | *p*-value | *z* | *p*-value |
| 0-10 |  |  |  |  |  |
| 10-20 | 65 | 3 | < .0001 | 0.4930 | 0.6220 |
| 20-30 | 4242 | 46 | < .0001 | 2.0680 | 0.0386 |
| >30 | 1401 | 26 | < .0001 | 3.3033 | 0.0010 |

| PT-PTS | Test of heterogeneity | | | Random effects model | |
| --- | --- | --- | --- | --- | --- |
|  | Q | n | *p*-value | *z* | *p*-value |
| 0-10 |  |  |  |  |  |
| 10-20 | 43 | 3 | < .0001 | 0.7252 | 0.4683 |
| 20-30 | 2650 | 19 | < .0001 | 1.3254 | 0.1851 |
| >30 | 333 | 16 | < .0001 | 3.2003 | 0.0014 |

| NT-NTS | Test of heterogeneity | | | Random effects model | |
| --- | --- | --- | --- | --- | --- |
|  | Q | n | *p*-value | *z* | *p*-value |
| 0-10 |  |  |  |  |  |
| 10-20 | 1049 | 5 | < .0001 | 1.3244 | 0.1854 |
| 20-30 | 3020 | 24 | < .0001 | 2.1644 | 0.0304 |
| >30 | 704 | 8 | < .0001 | -2.5294 | 0.0114 |

**Table S7.** Between-group heterogeneity (Qb) and probability (P) among n observations of carbon stock in response to different conservation agriculture practices under different sampling depth. For a description of the abbreviations, refer Fig. 2 title.

| PT-RT | Test of heterogeneity | | | Random effects model | |
| --- | --- | --- | --- | --- | --- |
|  | Q | n | *p*-value | *z* | *p*-value |
| Topsoil | 784 | 61 | < .0001 | 5.5514 | <.0001 |
| Subsoil | 401 | 38 | < .0001 | -2.9124 | 0.0036 |

| PT-NT | Test of heterogeneity | | | Random effects model | |
| --- | --- | --- | --- | --- | --- |
|  | Q | n | *p*-value | *z* | *p*-value |
| Topsoil | 33228 | 100 | < .0001 | 8.0485 | <.0001 |
| Subsoil | 1498 | 73 | < .0001 | -3.0433 | 0.0023 |

| RT-NT | Test of heterogeneity | | | Random effects model | |
| --- | --- | --- | --- | --- | --- |
|  | Q | n | *p*-value | *z* | *p*-value |
| Topsoil | 636 | 67 | < .0001 | 8.1547 | <.0001 |
| Subsoil | 142 | 34 | < .0001 | -1.2493 | 0.2116 |

| PTS-RTS | Test of heterogeneity | | | Random effects model | |
| --- | --- | --- | --- | --- | --- |
|  | Q | n | *p*-value | *z* | *p*-value |
| Topsoil | 120 | 33 | < .0001 | 1.1263 | 0.2600 |
| Subsoil | 168 | 15 | < .0001 | 2.0306 | 0.0423 |

| PTS-NTS | Test of heterogeneity | | | Random effects model | |
| --- | --- | --- | --- | --- | --- |
|  | Q | n | *p*-value | *z* | *p*-value |
| Topsoil | 1600 | 73 | < .0001 | 3.8942 | 0.0001 |
| Subsoil | 298 | 29 | < .0001 | 1.6574 | 0.0974 |

| RTS-NTS | Test of heterogeneity | | | Random effects model | |
| --- | --- | --- | --- | --- | --- |
|  | Q | n | *p*-value | *z* | *p*-value |
| Topsoil | 1502 | 24 | < .0001 | 1.9667 | 0.0492 |
| Subsoil | 160 | 16 | < .0001 | -0.1197 | 0.9047 |

| PT-RTS | Test of heterogeneity | | | Random effects model | |
| --- | --- | --- | --- | --- | --- |
|  | Q | n | *p*-value | *z* | *p*-value |
| Topsoil | 1681 | 21 | < .0001 | 4.7695 | <.0001 |
| Subsoil | 740 | 9 | < .0001 | -0.3470 | 0.7286 |

| PT-NTS | Test of heterogeneity | | | Random effects model | |
| --- | --- | --- | --- | --- | --- |
|  | Q | n | *p*-value | *z* | *p*-value |
| Topsoil | 3240 | 46 | < .0001 | 6.6410 | <.0001 |
| Subsoil | 2129 | 30 | < .0001 | -3.3380 | 0.0008 |

| PT-PTS | Test of heterogeneity | | | Random effects model | |
| --- | --- | --- | --- | --- | --- |
|  | Q | n | *p*-value | *z* | *p*-value |
| Topsoil | 2854 | 28 | < .0001 | 4.0826 | <.0001 |
| Subsoil | 954 | 12 | < .0001 | -1.8052 | 0.0710 |

| NT-NTS | Test of heterogeneity | | | Random effects model | |
| --- | --- | --- | --- | --- | --- |
|  | Q | n | *p*-value | *z* | *p*-value |
| Topsoil | 4435 | 25 | < .0001 | 4.4648 | <.0001 |
| Subsoil | 1148 | 13 | < .0001 | -0.9349 | 0.3498 |

**Figure captions**

**Fig. S1.** Boxplot with jitter plot of soil carbon stock to tillage practices under different agricultural tillage intensity (CAT: plow tillage, CT; reduced tillage, RT; or no tillage, NT), between residue removed and residue retained. Jitter shape indicates the cropping intensity, single, double and more (triple, or more than triple). Lower and upper whiskers are 25th and 75th percentiles, and lines within a box are medians.

**Fig. S2.** Relationships between mean annual precipitation (MAP) and soil C stock. Mean annual temperature (MAT) is indicated by color gradient.

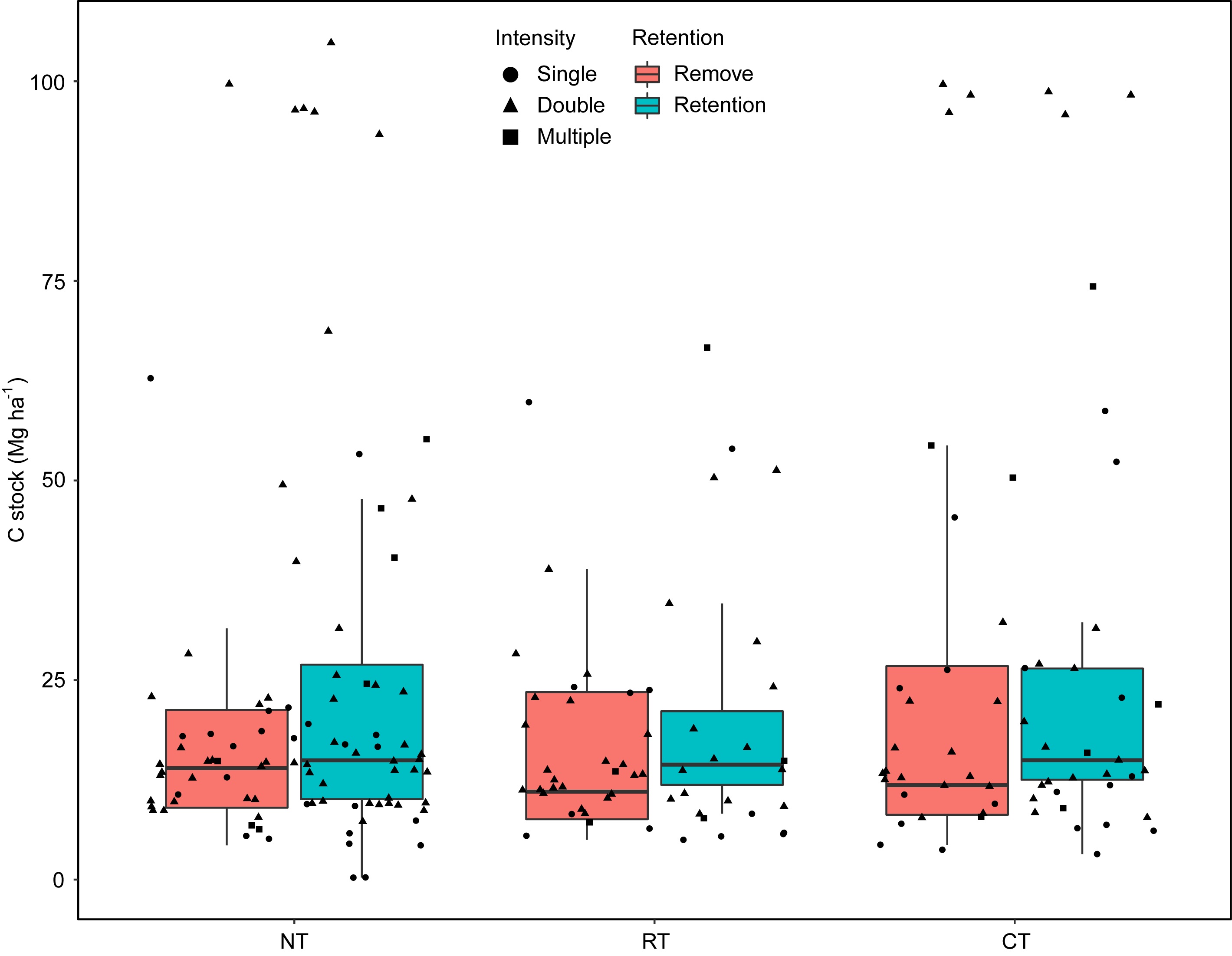

**Figure S1**

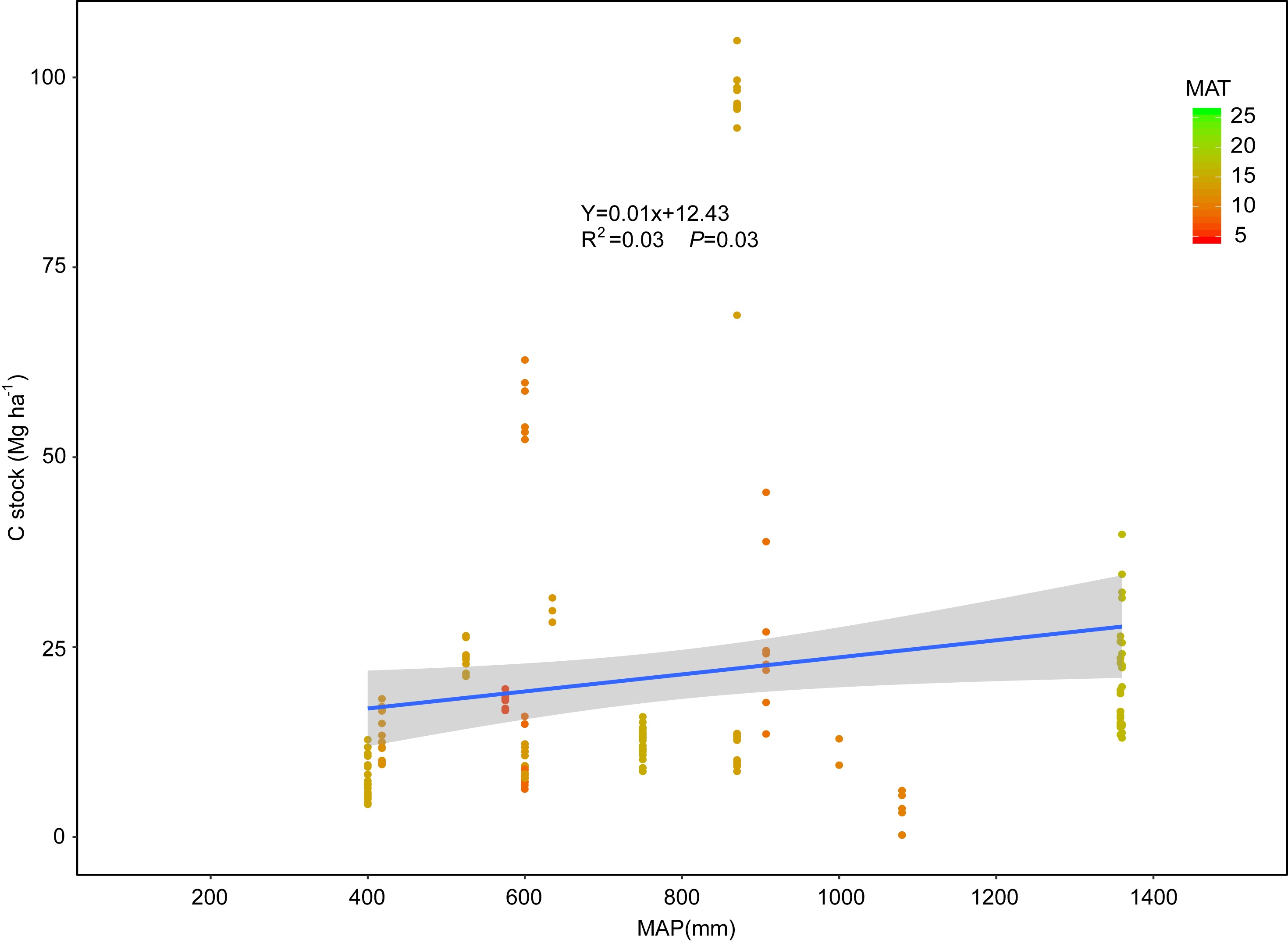

**Figure S2**

**Reference list for meta-analysis**

Abdollahi L, Getahun GT, Munkholm LJ (2017) Eleven years' effect of conservation practices for temperate sandy Loams: I. soil physical properties and topsoil carbon content. Soil Science Society of America Journal, 81, 380-391.

Abdollahi L, Munkholm LJ (2014) Tillage system and cover crop effects on soil quality: I. Chemical, mechanical, and biological properties. Soil Science Society of America Journal, 78, 262-270.

Afyuni M, Wagger MG (2006) Soil physical properties and bromide movement in relation to tillage system. Communications in soil science and plant analysis, 37, 541-556.

Alam M, Islam M, Salahin N, Hasanuzzaman M (2014) Effect of tillage practices on soil properties and crop productivity in wheat-mungbean-rice cropping system under subtropical climatic conditions. The Scientific World Journal, 2014, 1-15.

Álvaro-Fuentes J, Arrúe JL, Gracia R, López MV (2008) Tillage and cropping intensification effects on soil aggregation: Temporal dynamics and controlling factors under semiarid conditions. Geoderma, 145, 390-396.

Alvaro-Fuentes J, López MV, Cantero-Martínez C, Arrúe JL (2008) Tillage effects on soil organic carbon fractions in Mediterranean dryland agroecosystems. Soil Science Society of America Journal, 72, 541-547.

Apesteguía M, Virto I, Orcaray L, Bescansa P, Enrique A, Imaz MJ, Karlen DL (2017) Tillage effects on soil quality after three years of irrigation in northern Spain. Sustainability, 9, 1476.

Babujia LC, Hungria M, Franchini JC, Brookes PC (2010) Microbial biomass and activity at various soil depths in a Brazilian oxisol after two decades of no-tillage and conventional tillage. Soil Biology and Biochemistry, 42, 2174-2181.

Balota EL, Machineski O, Hamid KI, Yada IF, Barbosa GM, Nakatani AS, Coyne MS (2014) Soil microbial properties after long-term swine slurry application to conventional and no-tillage systems in Brazil. Science of the Total Environment, 490, 397-404.

Bescansa P, Imaz MJ, Virto I, Enrique A, Hoogmoed WB (2006) Soil water retention as affected by tillage and residue management in semiarid Spain. Soil and Tillage Research, 87, 19-27.

Bhattacharyya R, Tuti MD, Kundu S, Bisht JK, Bhatt JC (2012) Conservation tillage impacts on soil aggregation and carbon pools in a sandy clay loam soil of the Indian Himalayas. Soil Science Society of America Journal, 76, 617-627.

Bilalis D, Sidiras N, Vavoulidou E, Konstantas A (2009) Earthworm populations as affected by crop practices on clay loam soil in a Mediterranean climate. Acta Agriculturae Scandinavica Section B-Soil and Plant Science, 59, 440-446.

Bini D, dos Santos CA, Bernal LPT, Andrade G, Nogueira MA (2014) Identifying indicators of C and N cycling in a clayey Ultisol under different tillage and uses in winter. Applied Soil Ecology, 76, 95-101.

Blanco-Canqui H, Gantzer CJ, Anderson SH, Alberts EE (2004) Tillage and crop influences on physical properties for an Epiaqualf Soil Science Society of America Journal, 68, 567-576.

Blanco-Canqui H, Schlegel AJ, Heer WF (2011) Soil-profile distribution of carbon and associated properties in no-till along a precipitation gradient in the central Great Plains. Agriculture, Ecosystems & Environment, 144, 107-116.

Blanco-Canqui H, Stone LR, Schlegel AJ et al. (2009) No-till induced increase in organic carbon reduces maximum bulk density of soils. Soil Science Society of America Journal, 73, 1871-1879.

Blanco-Canqui H, Stone LR, Stahlman PW (2010) Soil response to long-term cropping systems on an Argiustoll in the central Great Plains. Soil Science Society of America Journal, 74, 602-611.

Blanco-Moure N, Gracia R, Bielsa AC, López MV (2016) Soil organic matter fractions as affected by tillage and soil texture under semiarid Mediterranean conditions. Soil and Tillage Research, 155, 381-389.

Bottinelli N, Hallaire V, Menasseri-Aubry S, Le Guillou C, Cluzeau D (2010) Abundance and stability of belowground earthworm casts influenced by tillage intensity and depth. Soil and Tillage Research, 106, 263-267.

Bravo CA, Giráldez JV, Ordóñez R, González P, Torres FP (2007) Long-term influence of conservation tillage on chemical properties of surface horizon and legume crops yield in a vertisol of southern Spain. Soil Science, 172, 141-148.

Büchi L, Wendling M, Amossé C, Jeangros B, Sinaj S, Charles R (2017) Long and short term changes in crop yield and soil properties induced by the reduction of soil tillage in a long term experiment in Switzerland. Soil and Tillage Research, 174, 120-129.

Carter MR (1992) Characterizing the soil physical condition in reduced tillage systems for winter wheat on a fine sandy loam using small cores. Canadian Journal of Soil Science, 72, 395-402.

Celik I, Günal H, Acar M, Gök M, Barut ZB, Pamiralan H (2017) Long-term tillage and residue management effect on soil compaction and nitrate leaching in a Typic Haploxerert soil. International Journal of Plant Production, 11, 131-150.

Chan KY, Heenan DP, Oates A (2002) Soil carbon fractions and relationship to soil quality under different tillage and stubble management. Soil and Tillage Research, 63, 133-139.

Chatterjee A, Lal R (2009) On farm assessment of tillage impact on soil carbon and associated soil quality parameters. Soil and Tillage Research, 104, 270-277.

Chen J, Pang DW, Han MM et al. (2017) Effects of tillage patterns on soil biological activity, availability of soil nutrients and grain yield of winter wheat Acta Agronomica Sinica, 43, 1245-1253.

Chen WC, Xu S, Zhu AN, Ma HW, He JQ, Liu JM (2015a) Effects of conservation tillage on the content of carbon, nitrogen in fluvo-aquic soil. Chinese Agricultural Science Bulletin, 31, 224-230.

Chen XW, Shi XH, Zhang XP, Liang AZ, Jia SX, Fan RQ, Wei SC (2011) No tillage impacts on soil organic carbon in a black soil in Northeast China. Fresenius Environmental Bulletin, 20, 3199-3205.

Chen ZD, Zhang HL, Dikgwatlhe SB, Xue JF, Qiu KC, Tang HM (2015b) Soil carbon storage and stratification under different tillage/residue-management practices in double rice cropping system. Journal of Integrative Agriculture, 14, 1551-1560.

Cheng C, Wang JJ, Cheng HH, Luo K, Zeng YJ, Shi QH, Shang QY (2018) Effects of straw returning and tillage system on crop yield and soil fertility quality in paddy field under double-cropping-rice system. Acta Pedologica Sinica, 55, 247-257.

Chowdhury S, Farrell M, Butler G, Bolan N (2015) Assessing the effect of crop residue removal on soil organic carbon storage and microbial activity in a no-till cropping system. Soil use and management, 31, 450-460.

Cui FJ, Liu JH, Li LJ, Gao J, Li Q (2012) Effect of zero tillage with mulching on active soil organic carbon. Acta Agriculturae Boreali-Occidentalis Sinica, 21, 195-200.

Dai XQ, Li YS, Ouyang Z, Wang HM, Wilson GV (2013) Organic manure as an alternative to crop residues for no-tillage wheat-maize systems in North China Plain. Field Crops Research, 149, 141-148.

Dai YX, Zhang XQ, Guo XX, Yang YM, Wang L, Liu JH (2016) Effects of no tillage with rotation on soil hydrothermal dynamics and available nutrients in dry farming area. Chinese Soil and Fertilization, 6, 14-20.

de Moraes Sa JC, Tivet F, Lal R, Briedis C, Hartman DC, dos Santos JZ, dos Santos JB (2014) Long-term tillage systems impacts on soil C dynamics, soil resilience and agronomic productivity of a Brazilian Oxisol. Soil and Tillage Research, 136, 38-50.

Dikgwatlhe SB, Chen ZD, Lal R, Zhang HL, Chen F (2014) Changes in soil organic carbon and nitrogen as affected by tillage and residue management under wheat-maize cropping system in the North China Plain. Soil and Tillage Research, 144, 110-118.

Dong W, Liu EK, Yan CR, Zhang HH, Zhang YQ (2017) Changes in the composition and diversity of topsoil bacterial, archaeal and fungal communities after 22 years conventional and no-tillage managements in Northern China. Archives of Agronomy and Soil Science, 63, 1369-1381.

Dong WX, Hu CS, Chen SY, Qin SP, Zhang YM (2013) Effect of conservation tillage on ammonia volatilization from nitrogen fertilizer in winter wheat-summer maize cropping system. Scientia Agricultura Sinica, 46, 2278-2284.

Duiker SW, Lal R (1999) Crop residue and tillage effects on carbon sequestration in a Luvisol in central Ohio. Soil and Tillage Research, 52, 73-81.

Fabrizzi KP, Moron A, García FO (2003) Soil carbon and nitrogen organic fractions in degraded vs. non-degraded Mollisols in Argentina. Soil Science Society of America Journal, 67, 1831-1841.

Fan RQ, Yang XM, Drury CF, Reynolds WD, Zhang XP (2014) Spatial distributions of soil chemical and physical properties prior to planting soybean in soil under ridge-, no-and conventional-tillage in a maize-soybean rotation. Soil use and management, 30, 414-422.

Feiziene D, Janusauskaite D, Feiza V, Putramentaite A, Sinkeviciene A, Suproniene S, Janusauskaite D (2015) After-effect of long-term soil management on soil respiration and otherqualitative parameters under prolonged dry soil conditions. Turkish Journal of Agriculture and Forestry, 39, 633-651.

Frey SD, Elliott ET, Paustian K (1999) Bacterial and fungal abundance and biomass in conventional and no-tillage agroecosystems along two climatic gradients. Soil Biology and Biochemistry, 31, 573-585.

Fuentes M, Govaerts B, De León F, Hidalgo C, Dendooven L, Sayre KD, Etchevers J (2009) Fourteen years of applying zero and conventional tillage, crop rotation and residue management systems and its effect on physical and chemical soil quality. European Journal of Agronomy, 30, 228-237.

Gajda AM, Czyż EA, Dexter AR, Furtak KM, Grządziel J, Stanek-Tarkowska J (2018) Effects of different soil management practices on soil properties and microbial diversity International Agrophysics, 32, 81-91.

Gajda AM, Czyż EA, Stanek-Tarkowska J, Dexter AR, Furtak KM, Grządziel J (2017) Effects of long-term tillage practices on the quality of soil under winter wheat. Plant, Soil and Environment, 63, 236-242.

Getahun GT, Munkholm LJ, Schjønning P (2016) The influence of clay-to-carbon ratio on soil physical properties in a humid sandy loam soil with contrasting tillage and residue management Geoderma, 264, 94-102.

Gil SV, Becker A, Oddino C, Zuza M, Marinelli A, March G (2009) Field trial assessment of biological, chemical, and physical responses of soil to tillage intensity, fertilization, and grazing. Environmental Management, 44, 378-386.

Gong WF, Li LL, Zhang XP, Shi DD (2013) Influence of conservation tllage on top soil physical and chemical quality in rain-fed areas of the Loess Plateau. Chinese Agricultural Science Bulletin, 29, 280-285.

Gosai K, Arunachalam A, Dutta BK (2010) Tillage effects on soil microbial biomass in a rainfed agricultural system of northeast India. Soil and Tillage Research, 109, 68-74.

Guo Q, Yu LL, Xu Y, Yang XH, Wang CL, Yang YF, Zhang SL (2015) Effects of conservation tillage on yield of corn and soil properties in the east of Hebei province. Journal of Hebei University of Science & Technology, 29, 14-17.

Han HF, Ning TY, Li ZJ, Cao HM (2014) Soil respiration rate in summer maize field under different soil tillage and straw application. Maydica, 59, 187-196.

Han HF, Ning TY, Li ZJ, Cao HM (2017) The ratio of CO_2_-Cemission to grain yield in summer maize cultivated under different soil tillage and straw application conditions. Experimental Agriculture, 53, 118-130.

He J, Li HW, Rasaily RG et al. (2011) Soil properties and crop yields after 11 years of no tillage farming in wheat-maize cropping system in North China Plain. Soil and Tillage Research, 113, 48-54.

He TB, Fan B, Li B et al. (2014) Effect of mechanized conservation tillage on soil physicochemical properties of dryland in Karst Mountainous area. Journal of Soil and Water Conservation, 28, 163-167.

He YY, Zhang HL, Sun GF, Tang WG, Li Y, Chen F (2010) Effect of different tillage on soil organic carbon and the organic carbon storage in two-crop paddy field. Journal of Agro-Environment Science, 29, 200-204.

Hermle S, Anken T, Leifeld J, Weisskopf P (2008) The effect of the tillage system on soil organic carbon content under moist, cold-temperate conditions. Soil and Tillage Research, 98, 94-105.

Himmelbauer ML, Sobotik M, Loiskandl W (2012) No-tillage farming, soil fertility and maize root growth. Archives of Agronomy and Soil Science, 58, S151-S157.

Hu N, Lou YL, Liang L (2009) Effects of the conservation tillage on soil nematode c-p groups and functional guilds. Ecology and Environmental Sciences, 18, 2349-2353.

Huang MX, Liang T, Wang LQ, Zhou CH (2015) Effects of no-tillage systems on soil physical properties and carbon sequestration under long-term wheat-maize double cropping system. Catena, 128, 195-202.

Hussain I, Olson KR, Ebelhar SA (1999) Long-term tillage effects on soil chemical properties and organic matter fractions. Soil Science Society of America Journal, 63, 1335-1341.

Jarecki MK, Lal R (2005) Soil organic carbon sequestration rates in two long-term no-till experiments in Ohio. Soil Science, 170, 280-291.

Jemai I, Aissa NB, Guirat SB, Ben-Hammouda M, Gallali T (2012) On-farm assessment of tillage impact on the vertical distribution of soil organic carbon and structural soil properties in a semiarid region in Tunisia. Journal of Environmental Management, 113, 488-494.

Ji Q, Sun HY, Taraqqi AK, Wang XD (2014) Impact of different tillage practices on soil organic carbon and water use efficiency under continuous wheat-maize binary cropping system. Chinese Journal of Applied Ecology, 25, 1029-1035.

Ji YY, Zhang GL, Zhang R, Liu YS, Yang DL (2012) Effects of different tillage modes on soil organic carbon and carbon pool management index in fluvo-aquic soils. Chinese Agricultural Science Bulletin, 28, 73-77.

Jia SX, Zhang XP, Chen XW et al. (2016) Long-term conservation tillage influences the soil microbial community and its contribution to soil CO_2_ emissions in a Mollisol in Northeast China. Journal of soils and sediments, 16, 1-12.

Jiang XB, Li YS, OUyang Z, Hou RX, Li FD (2012) Effect of no-tillage on soil aggregate and organic carbon storage Chinese Journal of Eco-Agriculture, 20, 270-278.

Karlen DL, Cambardella CA, Kovar JL, Colvin TS (2013) Soil quality response to long-term tillage and crop rotation practices. Soil and Tillage Research, 133, 54-64.

Kennedy AC, Schillinger WF (2006) Soil quality and water intake in traditional-till vs. no-till paired farms in Washington's Palouse region. Soil Science Society of America Journal, 70, 940-949.

Khan AU, Iqbal M, Islam KR (2007) Dairy manure and tillage effects on soil fertility and corn yields. Bioresource Technology, 98, 1972-1979.

Kibet LC, Blanco-Canqui H, Jasa P (2016) Long-term tillage impacts on soil organic matter components and related properties on a Typic Argiudoll. Soil and Tillage Research, 155, 78-84.

Kitur BK, Phillips SR, Olson KR, Ebelhar SA (1994) Tillage effects on selected chemical properties of Grantsburg silt loam. Communications in soil science and plant analysis, 25, 225-246.

Kumar S, Kadono A, Lal R, Dick W (2012) Long-term no-till impacts on organic carbon and properties of two contrasting soils and corn yields in Ohio. Soil Science Society of America Journal, 76, 1798-1809.

Lal R (1997) Soil degradative effects of slope length and tillage method on alfisols in western Nigeria II. Soil chemical properties, plant nutrient loss and water quality. Land Degradation & Development, 8, 221-244.

Lal R, Mahboubi AA, Fausey NR (1994) Long-term tillage and rotation effects on properties of a central Ohio soil. Soil Science Society of America Journal, 58, 517-522.

Li CF, Yue LX, Kou ZK, Zhang ZS, Wang JP, Cao CG (2012a) Short-term effects of conservation management practices on soil labile organic carbon fractions under a rape-rice rotation in central China. Soil and Tillage Research, 119, 31-37.

Li H, Zhang P, JIa ZK, Sun HX, Han LN, Yang BP, Nie JF (2012b) Effects of straw mulching treatment on characteristics of soil aggregates in Weibei dryland. Agricultural Research in the Arid Areas, 30, 27-33.

Li LJ, You MY, Shi HA, Han XZ (2012c) Short-term tillage influences microbial properties of a Mollisol in Northeast China. Journal of Food Agriculture & Environment, 10, 1433-1436.

Li T, Wang ZT, Liu L, Liao YC, Liu Y, Han J (2017) Effect of conservation tillage practices on soil microbial spatial distribution and soil physico-chemical properties of the northwest dryland. Scientia Agricultura Sinica, 50, 859-870.

Li WF, Liang AZ, Zhang XP, Shi XH, Shen Y, Fan RQ, Yang XM (2011) Short-term effects of no-tillage on soil organic C and N and available nutrients in black soil of Northeast China. Chinese Journal of Soil Science, 42, 664-669.

Li YP, Wang MB, Shi XY, Zhou J, Zhang XC (2012d) Influence of different tillage methods on soil physical and chemical properties and maize yield. Journal of Shanxi Agricultural Sciences, 40, 723-727.

Limousin G, Tessier D (2007) Effects of no-tillage on chemical gradients and topsoil acidification. Soil and Tillage Research, 92, 167-174.

Liu EK, Chen B, Yan C, Zhang Y, Mei X, Wang J (2015) Seasonal changes and vertical distributions of soil organic carbon pools under conventional and no-till practices on Loess Plateau in China. Soil Science Society of America Journal, 79, 517-526.

Liu EK, Teclemariam SG, Yan CR et al. (2014) Long-term effects of no-tillage management practice on soil organic carbon and its fractions in the northern China. Geoderma, 213, 379-384.

Liu EK, Zhao BQ, Mei XR, So HB, Li J, Li XY (2010) Effects of no-tillage management on soil biochemical characteristics in northern China. The Journal of Agricultural Science, 148, 217-223.

Liu PT, Feng BL, Mu F et al. (2009) Effects of conservation tillage on soil physicochemical properties in the spring maize area of the Loess Plateau. Agricultural Research in the Arid Areas, 27, 171-175.

Liu XM, Li Q, Liang WJ, Jiang Y, Wen DZ (2006) Dynamics of aquicbrown soil enzyme activities under no-tillage. Chinese Journal of Applied Ecology, 17, 1347-1351.

López-Fando C, Dorado J, Pardo MT (2007) Effects of zone-tillage in rotation with no-tillage on soil properties and crop yields in a semi-arid soil from central Spain. Soil and Tillage Research, 95, 266-276.

López-Fando C, Pardo MT (2009) Changes in soil chemical characteristics with different tillage practices in a semi-arid environment. Soil and Tillage Research, 104, 278-284.

López-Garrido R, Deurer M, Madejón E, Murillo JM, Moreno F (2012) Tillage influence on biophysical soil properties: The example of a long-term tillage experiment under Mediterranean rainfed conditions in South Spain. Soil and Tillage Research, 118, 52-60.

Lv RZ, Huang M, Xiong Y, Li YJ, Zhang J, Sun HZ (2015) Effect of tillage methods on soil physico-chemical properties and enzymes activity under pea-wheat rotation. Jiangsu Journal of Agricultural Science, 43, 100-103.

Lv RZ, Xiong Y, Li YJ, Li Q, Huang M (2014) Effect ofconservation tillage on soil carbon pool in farmland. Journal of Soil and Water Conservation, 28, 206-217.

Lv YZ, Lian XJ, Zhao H, Liu WR (2010) Effects of conservation tillage patterns on content and density of organic carbon of black soil. Transactions of the CSAE, 26, 163-169.

Mahboubi AA, Lal R, Faussey NR (1993) Twenty-eight years of tillage effects on two soils in Ohio. Soil Science Society of America Journal, 57, 506-512.

Małecka I, Blecharczyk A, Sawinska Z, Swędrzyńska D, Piechota T (2015) Winter wheat yield and soil properties response to long-term non-inversion tillage. Journal of Agricultural Science and Technology, 17, 1571-1584.

Martín-Lammerding D, Navas M, del Mar Albarrán M, Tenorio JL, Walter I (2015) LONG term management systems under semiarid conditions: Influence on labile organic matter, β-glucosidase activity and microbial efficiency. Applied Soil Ecology, 96, 296-305.

Mazzoncini M, Antichi D, Di Bene C, Risaliti R, Petri M, Bonari E (2016) Soil carbon and nitrogen changes after 28 years of no-tillage management under Mediterranean conditions. European Journal of Agronomy, 77, 156-165.

McVay KA, Budde JA, Fabrizzi K et al. (2006) Management effects on soil physical properties in long-term tillage studies in Kansas. Soil Science Society of America Journal, 70, 434-438.

Melero S, Panettieri M, Madejón E, Macpherson HG, Moreno F, Murillo JM (2011) Implementation of chiselling and mouldboard ploughing in soil after 8 years of no-till management in SW, Spain: Effect on soil quality. Soil and Tillage Research, 112, 107-113.

Melero S, Vanderlinden K, Ruiz JC, Madejón E (2009) Soil biochemical response after 23 years of direct drilling under a dryland agriculture system in southwest Spain. The Journal of Agricultural Science, 147, 9-15.

Moebius-Clune BN, van Es HM, Idowu OJ et al. (2008) Long-term effects of harvesting maize stover and tillage on soil quality. Soil Science Society of America Journal, 72, 960-969.

Motta AC, Reeves DW, Touchton JT (2002) Tillage intensity effects on chemical indicators of soil quality in two coastal plain soils. Communications in soil science and plant analysis, 33, 913-932.

Mukherjee A, Lal R (2015) Tillage effects on quality of organic and mineral soils under on-farm conditions in Ohio. Environmental Earth Sciences, 74, 1815-1822.

Muñoz A, López-Piñeiro A, Ramírez M (2007) Soil quality attributes of conservation management regimes in a semi-arid region of south western Spain. Soil and Tillage Research, 95, 255-265.

Naab JB, Mahama GY, Yahaya I, Prasad PVV (2017) Conservation agriculture improves soil quality, crop yield, and incomes of smallholder farmers in north western Ghana. Frontiers in Plant Science, 8, 1-18.

Niu YN, Zhang RZ, Luo ZZ, Li LL, Cai LQ, Li G, Xie JH (2016) Contributions of long-term tillage systems on crop production and soil properties in the semi-arid Loess Plateau of China. Journal of the Science of Food and Agriculture, 96, 2650-2659.

Nugis E, Edesi L, Tamm K, KADAJA J, Akk E, Viil P, ILUMÄE E (2016) Response of soil physical properties and dehydrogenase activity to contrasting tillage systems. Zemdirbyste-Agriculture, 103, 123-128.

Okeyo JM, Norton J, Koala S, Waswa B, Kihara J, Bationo A (2016) Impact of reduced tillage and crop residue management on soil properties and crop yields in a long-term trial in western Kenya. Soil Research, 54, 719-729.

Oorts K, Garnier P, Findeling A, Mary B, Richard G, Nicolardot B (2007) Modeling soil carbon and nitrogen dynamics in no-till and conventional tillage using PASTIS model. Soil Science Society of America Journal, 71, 336-346.

Pan YW, Fan J, Hao MD, Chen X (2016) Effects of long-term tillage and mulching methods on properties of surface soil and maize yield in tableland region of the Loess Plateau. Journal of Plant Nutrition and Fertilizer, 22, 1558-1567.

Pang X, He WQ, Yan CR, Liu EK, Liu S, Yin T (2013) Effect of tillage and residue management on dynamic of soil microbial biomass carbon. Acta Ecologica Sinica, 33, 1308-1316.

Pei XX, Dang JY, Zhang DY, Wang JA, Zhang J (2014) Effects of different tillage methods on phospholipid fatty acids and enzyme activities in calcareous cinnamon soil. Chinese Journal of Applied Ecology, 25, 2275-2280.

Qin SP, He XH, Hu CS, Zhang YM, Dong WX (2010a) Responses of soil chemical and microbial indicators to conservational tillage versus traditional tillage in the North China Plain. European Journal of Soil Biology, 46, 243-247.

Qin SP, Hu CS, Wang YY, Li XX, He XH (2010b) Tillage effects on intracellular and extracellular soil urease activities determined by an improved chloroform fumigation method. Soil Science, 175, 568-572.

Roldán A, Caravaca F, Hernández MT, Garcıa C, Sánchez-Brito C, Velásquez M, Tiscareno M (2003) No-tillage, crop residue additions, and legume cover cropping effects on soil quality characteristics under maize in Patzcuaro watershed (Mexico). Soil and Tillage Research, 72, 65-73.

Saha S, Chakraborty D, Sharma AR, Tomar RK, Bhadraray S, Sen U, Kalra N (2008) Effect of tillage and residue management on soil physical properties and crop productivity in maize (*Zea mays*)-Indian mustard (*Brassica juncea*) system. Indian Journal of Agricultural Sciences, 80, 679-685.

Sasal MC, Andriulo AE, Taboada MA (2006) Soil porosity characteristics and water movement under zero tillage in silty soils in Argentinian Pampas. Soil and Tillage Research, 87, 9-18.

Schjønning P, Thomsen IK, Petersen SO, Kristensen K, Christensen BT (2011) Relating soil microbial activity to water content and tillage-induced differences in soil structure. Geoderma, 163, 256-264.

Schmidt ES, Villamil MB, Amiotti NM (2018) Soil quality under conservation practices on farm operations of the southern semiarid pampas region of Argentina. Soil and Tillage Research, 176, 85-94.

Shao YH, Xie YX, Wang CY et al. (2016) Effects of different soil conservation tillage approaches on soil nutrients, water use and wheat-maize yield in rainfed dry-land regions of North China. European Journal of Agronomy, 81, 37-45.

Sharma KL, Grace JK, Srinivas K et al. (2009) Influence of tillage and nutrient sources on yield sustainability and soil quality under sorghum-mung bean system in rainfed semi-arid tropics. Communications in soil science and plant analysis, 40, 2579-2602.

Sharma KL, Mandal UK, Srinivas K, Vittal KPR, Mandal B, Grace JK, Ramesh V (2005) Long-term soil management effects on crop yields and soil quality in a dryland Alfisol. Soil and Tillage Research, 83, 246-259.

Shukla MK, Lal R, Ebinger M (2003) Tillage effects on physical and hydrological properties of a typic Argiaquoll in central Ohio. Soil Science, 168, 802-811.

Spedding TA, Hamel C, Mehuys GR, Madramootoo CA (2004) Soil microbial dynamics in maize-growing soil under different tillage and residue management systems. Soil Biology and Biochemistry, 36, 499-512.

Su L, Zhang RZ, Cai LQ (2012) Effects of conservation tillage on organic carbon content in soil. Jiangsu Journal of Agricultural Science, 28, 524-529.

Su LL, Li YJ, Xu WX, Tang JH, Chen CX, Hao WW, Wang N (2017) Effects of tillage methods on soil physical and chemical properties and yield of summer soybean. Agricultural Research in the Arid Areas, 35, 43-48.

Sui PX, Zhang XY, Wen XF, You DB, Tian P, Qi H (2016) Effects of tillage and straw management on nutrient contents and enzyme activities of brown soil. Chinese Journal of Ecology, 35, 2038-2045.

Sun B, Jia SX, Zhang SX, McLaughlin NB, Liang AZ, Chen XW, Zhang XP (2016) No tillage combined with crop rotation improves soil microbial community composition and metabolic activity. Environmental Science and Pollution Research, 23, 6472-6482.

Sun J, Liu M, Li LJ, Liu JH, Acharya SN (2010) Effects of different tillage systems on soil hydrothermal regimes in rain-fed field of Inner Mongolia. Acta Ecologica Sinica, 30, 1539-1547.

Sweeney DW (2017) Does 20 Years of Tillage and N Fertilization Influence Properties of a Claypan Soil in the Eastern Great Plains? Agricultural & Environmental Letters, 2, 1-4.

Tarkalson DD, Hergert GW, Cassman KG (2006) Long-term effects of tillage on soil chemical properties and grain yields of a dryland winter wheat-sorghum/corn-fallow rotation in the Great Plains. Agronomy journal, 98, 26-33.

Wander MM, Bidart MG, Aref S (1998) Tillage impacts on depth distribution of total and particulate organic matter in three Illinois soils. Soil Science Society of America Journal, 62, 1704-1711.

Wang CX, Wang XD, Zhu RX (2011a) Effect of conservational tillage measures on the oxidation stability of soil organic carbon in Soil aggregates. Chinese Agricultural Science Bulletin, 26, 121-126.

Wang DD, Zhou L, Huang SQ, Li CF, Cao CG (2013a) Short-term effects of tillage practices and wheat straw returned to the field on topsoil labile organic carbon fractions and yields in central China. Journal of Agro-Environment Science, 32, 735-740.

Wang GL, Hao MD, Xu JG, Hong JP (2011b) Effect of conservation tillage on wheat yield and soil physicochemical properties in the south of Loess Plateau. Plant Nutrition and Fertilizer Science, 17, 539-544.

Wang HF, Zhang YQ, Wu ZH, Zhang JC, Qiao SS (2011c) Effects of long-term zero-tillage on soil physiochemical properties and enzyme activities in cinnamon soil. Agricultural Research in the Arid Areas, 29, 136-141.

Wang L, Li LL, Gao LF, Liu J, Luo ZZ, Xie JH (2013b) Effect of long-term conservation tillage on total organic carbon and readily oxidizable organic carbon in loess soils. Chinese Journal of Eco-Agriculture, 21, 1057-1063.

Wang QJ, Bai YH, Gao HW et al. (2008) Soil chemical properties and microbial biomass after 16 years of no-tillage farming on the Loess Plateau, China. Geoderma, 144, 502-508.

Wang RL, Zhang ZD, LIu JH, Liu HJ, Liu JQ, Wu JY (2016a) Effects of no-tillage with mulching on soil nutrient content and microbial biomass in oat field. Journal of Soil and Water Conservation, 30, 183-193.

Wang Z, Liu L, Chen Q, Wen X, Liao Y (2016b) Conservation tillage increases soil bacterial diversity in the dryland of northern China. Agronomy for Sustainable Development, 36, 28.

Wu JF, Zeng YH, Zhao XF, Fan CG, Pan XH, Shi QH (2017) Effects of tillage methods on yield of double cropping rice with machine-transplanted and soil physical-chemical properties. Journal of Hunan Agricultural University (Natural Sciences), 43, 581-585.

Wu YH, Tiao XH, Chi WB, Nan XX, Yan XL, Zhu RX, Tong YA (2010) Numerical evaluation of soil quality under different conservation tillage patterns. Chinese Journal of Applied Ecology, 21, 1468-1476.

Wyngaard N, Echeverría HE, Rozas HRS, Divito GA (2012) Fertilization and tillage effects on soil properties and maize yield in a Southern Pampas Argiudoll. Soil and Tillage Research, 119, 22-30.

Xu SQ, Zhang RZ, Dong B, Zhang M (2009) Effect of tillage practices on structural properties and content of organic carbon in tilth soil. Chinese Journal of Eco-Agriculture, 17, 203-208.

Xue JF, Pu C, Liu SL, Chen ZD, Chen F, Xiao XP, Zhang HL (2015) Effects of tillage systems on soil organic carbon and total nitrogen in a double paddy cropping system in Southern China. Soil and Tillage Research, 153, 161-168.

Yan CR, Liu EK, He WQ, Liu S, Liu Q (2010) Effect of different tillage on soil organic carbon and its fractions in the Loess Plateau of China. Chinese Soil and Fertilization, 6, 58-63.

Yan J, Deng LJ, Huang J (2005) Effeet of conservaiton tillage on soil physicochemieal properites and crop yields Chinese Agricultural Mechanzation, 2, 31-34.

Yang AN, Hu JH, Lin XG, Zhu AN, Wang JH, Dai J, Wong MH (2012) Arbuscular mycorrhizal fungal community structure and diversity in response to 3-year conservation tillage management in a sandy loam soil in North China. Journal of soils and sediments, 12, 835-843.

Yao SL, Teng XL, Zhang B (2015) Effects of rice straw incorporation and tillage depth on soil puddlability and mechanical properties during rice growth period. Soil and Tillage Research, 146, 125-132.

Yin ZY, Huang L, Xue B, Huang YN, Li XK, Lu JW (2017) Effect of conservation tillage on soil fertility under rice-rape rotation system. Chinese Journal of Eco-Agriculture, 25, 1604-1614

You D, Tian P, Sui P, Zhang W, Yang B, Qi H (2017) Short-term effects of tillage and residue on spring maize yield through regulating root-shoot ratio in Northeast China. Scientific reports, 7, 13314.

Yu HY, Peng WY, Ma X, Zhang KL (2011) Effects of no-tillage on soil water content and physical properties of spring corn fields in semi-arid region of northern China. Chinese Journal of Applied Ecology, 22, 99-104.

Zhang B, He HB, Ding XL, Zhang XD, Zhang XP, Yang XM, Filley TR (2012) Soil microbial community dynamics over a maize (*Zea mays* L.) growing season under conventional-and no-tillage practices in a rainfed agroecosystem. Soil and Tillage Research, 124, 153-160.

Zhang J, Yao YQ, Lu JJ et al. (2008) Soil carbon change and yield increase mechanism of conservation tillage on sloping drylands in semi-humid arid area. Chinese Journal of Eco-Agriculture, 16, 297-301.

Zhang J, Zhang RZ, Zuo XA (2016a) Effect of conservation tillage on physical and chemical characteristics under pea-wheat rotation system in the Loess Plateau. Journal of Desert Research, 36, 137-143.

Zhang L, Yao YQ, Jin K et al. (2007) Change of MBC and MBN under conservation tillage on sloping dryland. Journal of Soil and Water Conservation, 21, 126-129.

Zhang M, Li LK, Hao MD (2013) Effect of no-tillage with straw cover on corn yield and soil fertility. Acta Agriculturae Boreali-Occidentalis Sinica, 22, 67-72.

Zhang MY, Huang GH, Kong FL, Chen F, Zhang HL (2011) Influences of tillage practices on distribution of soil microbial biomass carbon under winter wheat in north China Ecology and Environmental Sciences, 20, 409-414.

Zhang X, Li H, He J, Wang Q, Golabi MH (2009a) Influence of conservation tillage practices on soil properties and crop yields for maize and wheat cultivation in Beijing, China. Soil Research, 47, 362-371.

Zhang XJ, Liu JH, Li LJ, Duan LK, Wang ZG, Su SH (2009b) Effects of different conservation tillage on soil microbes quantities and enzyme activities in dry cultivation. Chinese Journal of Soil Science, 40, 542-546.

Zhang ZQ, Qiang HJ, McHugh AD, He J, Li HW, Wang QJ, Lu ZY (2016b) Effect of conservation farming practices on soil organic matter and stratification in a mono-cropping system of Northern China. Soil and Tillage Research, 156, 173-181.

Zhao RL, Liu PT, Feng BL et al. (2010) Effect of different cultivation measures on activity and bound forms of organic carbon in Lou soil. Agricultural Research in the Arid Areas, 28, 69-74.

Zhou H, Lv YZ, Yang ZC, Li BG (2007) Effects of conservation tillage on soil aggregates in Huabei Plain, China Scientia Agricultura Sinica, 40, 1973-1979.

Zhu L, Hu N, Yang M, Zhan X, Zhang Z (2014) Effects of different tillage and straw return on soil organic carbon in a rice-wheat rotation system. PloS one, 9, e88900.
